# Diet induced mitochondrial DNA replication instability in *Rad51c* mutant mice drives sex-bias in anemia of inflammation

**DOI:** 10.1101/2024.09.21.613572

**Authors:** Karl-Heinz Tomaszowski, Yue Chen, Sunetra Roy, Momo Harris, Jianhua Zhang, Chi-lin Tsai, Katharina Schlacher

## Abstract

Anemia of inflammation (AI) is a common comorbidity associated with obesity, diabetes, cardiac disease, aging, and during anti-cancer therapies. Mounting evidence illustrates that males are disproportionally affected by AI, but not why. Here we demonstrate a molecular cause for a sex-bias in inflammation. The data shows that mitochondrial DNA (mtDNA) instability induced by dietary stress causes anemia associated with inflamed macrophages and improper iron recycling in mice. These phenotypes are enhanced in mice with mutations in *Fanco/Rad51c*, which predisposes to the progeroid disease Fanconi Anemia. The data reveals a striking sex-bias whereby females are protected. We find that estrogen acts as a mitochondrial antioxidant that reduces diet-induced oxidative stress, mtDNA replication instability and the distinctively mtDNA-dependent unphosphorylated STAT1 response. Consequently, treatment of male *Rad51c* mutant mice with estrogen or mitochondrial antioxidants suppresses the inflammation-induced anemia. Collectively, this study uncovers estrogen-responsive mtDNA replication instability as a cause for sex-specific inflammatory responses and molecular driver for AI.

## Background

Unhealthy diets are prominent sources of environmental stress that are major contributors to the etiology of many diseases. Many components of a modern Western diet including high fat intake, mitogenic toxins from over-charred meats and excessive alcohol consumption potently contribute to obesity, diabetes, heart attack, immune diseases, cancer and accelerate aging in the general population ^1^. Underlying genetic predisposition can further exacerbate disease severity, shortening onset of disease and accelerating aging processes. Amongst others, heterozygous mutations in Fanconi anemia (FA) and breast cancer (BRCA) disease susceptibility genes increase the risk to diabetes, cardiovascular disease, cancer and accelerate aging ^2–6^.

Many of the dietary agents are DNA-reactive and damage DNA; obesity has been shown to increase gH2AX ^7^, a molecular marker for DNA replication stress and DNA breaks, over-charred foods results in polycyclic aromatic hydrocarbons (PAH) that intercalate with DNA ^8–10^, and acetaldehydes from alcohol consumption create toxic DNA and protein-DNA crosslinks ^11^. Additionally, nutrition agents cause DNA-reactive oxygen species (ROS) ^12^. Any of these DNA damages in turn potently blocks DNA replication, jeopardizing genome stability. FA/BRCA genes play a critical role in ensuring the stability of DNA replication forks, in particular at the level of fork protection ^13,14^, which is a genome stability pathway that suppresses classic cGAS-dependent inflammation resulting in Type I interferons IFNa and IFNb ^15–18^.

In addition to nuclear DNA, environmental DNA-reactive stresses also damage mitochondrial DNA. FANC/BRCA genes have a dual role and act also in mitochondria in the proper clearance of damaged mitochondria ^19^, and at the level of mtDNA replication stability ^20^. Specifically, FA/BRCA promote mtDNA stability by preventing the degradation of nascently replicated mtDNA ^20^. It is known that mtDNA instability too elicits a cGAS-dependent inflammation response ^20^. However, it is a distinct inflammatory signaling termed the unphosphorylated STAT1 (un-P-STAT1) response. This response differs from the canonical cGAS activation in that it does not involve STAT1 phosphorylation at tyrosine 701, which is required to stimulate IFNa/b transcription, nor JAK pathway activation ^21–23^. Instead, total STAT1 protein is increased, resulting in a distinct interferon-stimulated-gene (ISG) expression signature that upregulates ∼30 of the over 300 gene products of the type I IFN response pathway ^20–26^.

One of the most common co-morbidities of diet-accelerated diseases including obesity, diabetes, cardiac disease and aging in the general population is anemia of chronic disease, also termed anemia of inflammation (AI) ^27–29^. AI also is the most frequent anemia seen in hospitalized and chronically ill patients, and one of the most prevalent side-effects during cancer and of anti-cancer therapies ^27–30^.

In contrast to iron-deficient anemia, AI is characterized by increased iron stores caused by inflamed and reprogrammed macrophages in the spleen, liver and bone marrow that are no longer able to properly recycle iron from erythrocytes, thereby curtailing erythropoiesis. While the direct role of inflammation in the progression of the disease remains elusive, it is known to involve rewired cytokine signaling involving Il-6, IFNg, Il-1 and TNFa. This in turn activates hepcidin transcription, which is the central iron metabolism-regulating hormone hepcidin. Hepcidin inhibits the Fe2+ exporter ferroportin in macrophages and so the cellular export of iron leading to the retention of iron within macrophages. Instead, bio-inactive Fe3+ is stored within the hemosiderin protein complex, which causes a granular macrophage appearance, and eventually iron release into the surrounding tissue ^27–29^. Thus, in contrast to iron-deficient anemia, which lacks the iron required during erythropoiesis, AI is not caused by a deficiency of iron per so, but rather by the inability of inflamed macrophages to properly metabolize it.

Mounting evidence suggest an unresolved gender bias seen in AI, whereby males show a collective increased risk for developing AI in conjunction with aging and in patients with diabetes or cardiac disease ^31–38^, pointing towards sex-specific inflammation programs. The underlying cause for this specific disparity between males and female predisposition is currently unknown. Here we identify mitochondrial replication instability as a key molecular driver for sex specific differences in inflammatory responses driving anemia of inflammation.

## Results

### Chronic exposure to Western-like diet causes male-specific anemia

Both genetic predisposition and exposure to environmental factors can contribute to human disease. Modern Western diets have changed significantly over the past century ^11^. Concomitant with an increase in the amounts of fat, sugar, protein and dietary carcinogens, is an increase in the risk of diabetes, cardiac disease, cancer and accelerated aging ^11^. To better understand the effects of chronic exposure to a modern Western diet, we exposed mice to a high-fat diet (HFD, 60% fat) and weekly treatments with 7,12-dimethylbenz(a)anthracene (DMBA), a carcinogenic polycyclic aromatic hydrocarbon similar to those found in over-charred meats and proteins ^8–10^ (Fig. 1A). As genetic factors can accelerate the effects of diet induced diseases, we included both wild-type and *FancO/Rad51c^dah/dah^*mutant mice. We focused on *FancO/Rad51c* mice as Fanconi Anemia (FA) is a genetic disease encompassing pleiotropic phenotypes including accelerated aging, cancer, bone marrow failure, metabolic disease, increased risk of cardiac arrest, and increased inflammation ^39–41^. Thus, mutations in FA genes predispose to many of the diseases that are caused by adverse diet. Moreover, heterozygous FA/BRCA mutations have been implicated in predisposing to environmental carcinogen induced diseases, including cancer and diabetes ^42,43^. Importantly, FA mutations are widely prevalent in the general western population, with 75% of the population harboring heterozygous disease-associated mutations in a FANC gene, making FA a relevant genetic model for the general population ^44^.

**Fig. 1.**
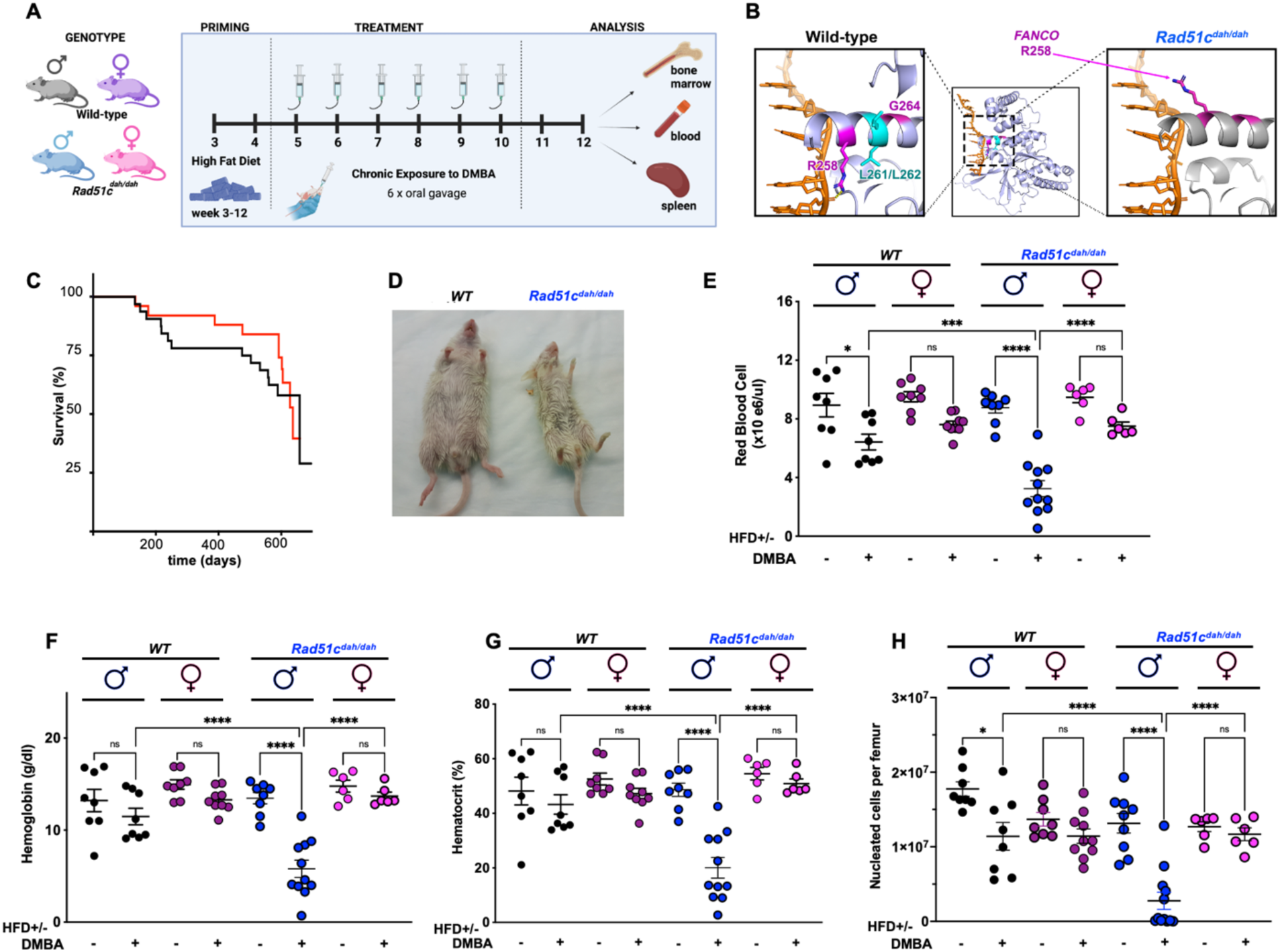
Male mice are prone to anemia under chronic stress exposure. **A)** Experimental workflow for mouse experiments under chronic stress treatment. **B)** Structure model using AF3 of wild-type (WT) and mutant Rad51c^dah/dah^ on DNA. Left: Magnification of WT Rad51c depicting murine R267 (human FA mutations R258H) in pink, which interacts with DNA (green) in this projection. Rad51c L270/L271 depicted in yellow, are deleted in the Rad51c^dah/dah^ mouse (zoom to the right). The deletion is predicted to functionally impacts R267 by changing the rotation of the alpha helix and **C)** Kaplan-Meyer survival curve of Rad51c WT and Rad51c^dah/dah^ mice. **D)** Representative picture of high fat diet and 7,12-dimethylbenz(a)anthracene (DMBA)-treated male Rad51 WT (left) and anemic male Rad51c dah/dah mice (right). **E)** Scatter dot plot of red cell blood count from chronic stress experiments. Male WT mice are shown in black, female WT mice are shown in purple, male Rad51c^dah/dah^ mice are shown in blue and female Rad51c^dah/dah^ mice are shown in pink **F-H)** Scatter dot plot of hemoglobin count (**F**), hematocrit (**G**) and nucleated cells per femur (**H**) from chronic stress experiments. Horizontal bars represent the mean and error bars represent the standard error of the mean. p-values are derived using the one-way ANOVA test. ns p>0.05, * p<0.05, **p < 0.01, ***p < 0.001, ****p<0.0001, n=6-11

We have created a CRISPR/Cas9 mutant mouse model missing two-amino-acids (*Rad51c^dah/dah^*for “**d**eletion in **a**lpha **h**elix” ^45^,missing L270/L271, corresponding to human L261/L262) in *Fanco/Rad51c*. This deletion is located between two FA patient associated mutation sites (human R258H and G264S) ^45^ ^46^. These alterations are part of a functional alpha-helix that influences a key DNA interaction motif loop ^47^. Using Alphafold 3 in comparison with recently solved crystal structures of RAD51C ^47^, we modeled the consequences of the deletion mutation in our mouse model (Fig. 1B). The projection suggests that the sidechain at position 258 interacts with DNA in the wild-type protein, but points away from DNA with the helical rotation caused by the two-amino acid deletion in our mouse mutant protein, suggesting functional impacts (Fig. 1B, right model). Yet, the homozygous hypomorphic *Rad51c^dah/dah^* mutant mice kept under unchallenged conditions in a pure FEVB strain background develop normally, are born with an expected Mendelian frequency and show no adverse phenotype on overall survival or body weight with standard CHOW feed under unchallenged conditions (Figs. 1C, S1A and Table S2), making it an optimal model system to test genetic potentiation.

To test the impact of modern Western diet components, both wild-type (WT) FVB mice and *Rad51c^dah/dah^*mutant mice were fed with a HFD after weaning and injected with DMBA weekly starting at 5 weeks of age (.1A). We noticed that male DMBA-treated *Rad51c^dah/dah^* mice developed weight loss and discoloration of paws within two weeks after treatment (Fig. 1D), both characteristic symptoms of anemia in mice. Blood analysis (Figs. 1E-1G) revealed that male WT mice treated with both HFD+DMBA showed a significant reduction in red blood cells (RBCs, Fig. 1E). Consistent with a role for genetic variation as an exacerbating factor, male *Rad51c^dah/dah^* mutant mice showed greater reduction in RBCs compared with WT mice (Figs. 1D). Similar trends were observed in hemoglobin and hemocrit levels (Figs 1F and 1G), as markers of anemia. Interestingly, when feeding the mice a diet with 10% instead of 60% fat, the DMBA treatment did cause a reduction in red blood cell values (Figs. S1B-S1D), but to a much lesser extent. This suggests that a HFD accelerates the anemia process with the combination of factors resulting in severe anemia. Strikingly, female mice are protected from the treatment induced anemia regardless of *Rad51c* status (Fig. 1D-1G).

As the bone marrow tissue is the origin of hematopoiesis ^48^, we analyzed the number of nucleated bone marrow cells (Fig. 1H). The data shows a strong reduction in nucleated bone marrow cells in the anemic male *Rad51c ^dah/dah^* mice (Fig. 1H). Similar to RBC’s, WT males too show a significant, albeit less pronounced, reduction in total bone marrow cells, while females of both genotypes are protected and asymptomatic (Fig. 1H). As bone marrow contains hematopoietic stem cells and progenitor cells of all blood cell lineages, we further analyzed white blood cell (WBC) and platelet counts (Fig. S1E and S1F). This blood analysis revealed similar trends in WBC and platelet count reductions as seen for the RBCs, with male *Rad51c* mice showing the most pronounced decrease in cell counts (Fig. S1E and S1F). While anemia is the most prominent phenotype, these analyses suggest additional milder pancytopenic effects.

Taken together, these data indicate that mice develop anemia under chronic stress conditions as a primary effect. In contrast to classical iron-deficiency anemia, this phenotype appears to preferentially occur in male mice.

### Male-specific anemia is associated with inflammation

To gain insights into how the male mice develop anemia, we performed bulk RNA-sequencing analysis in bone marrow of acutely DMBA-treated mice (Fig. 2A). Consistent with the overall phenotype, the largest difference in differentially expressed genes (DEG) between the groups is detected between male and female *Rad51c^dah/dah^* mice, confirming the phenotype-amplifying effect of the *Rad51c* mutation (Fig. 2B). The largest changes in the male mice were detected in upregulated genes (Fig. 2B and 2C). This is similar to the results when comparing DMBA-treated to non-treated mice (Fig. S2A), indicating that DMBA predominantly induces gene expression under these conditions.

**Fig. 2.**
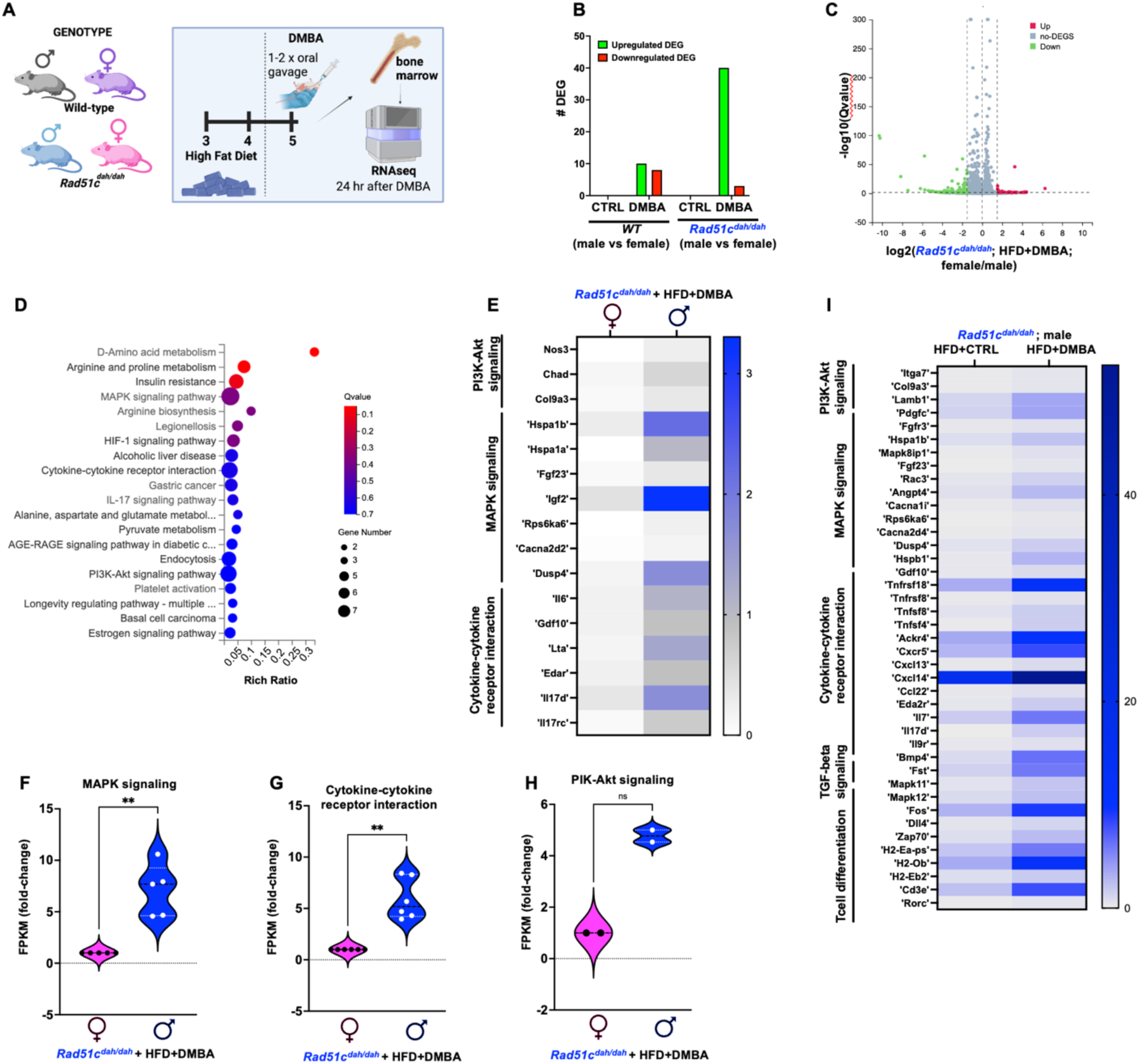
Dietary stress induces inflammation in male *Rad51c^dah/dah^* mice. **A)** Experimental workflow for short-term treatments. **B)** Number of differential expressed genes (DEG) in male compared to female WT and *Rad51c^dah/dah^* bone marrow. CTRL, control. All animals are fed with high fat diet (HFD). **C)** Volcano plot of up- and downregulated genes in male compared to female *Rad51c^dah/dah^* mice. **D)** KEGG pathway analysis of upregulated genes from DMBA-treated male compared to female *Rad51c^dah/dah^* mice, highlighting upregulation of inflammation pathways. **E)** RNAseq data heatmap of inflammatory pathways from KEGG analysis in (**D**) comparing male and female HFD+DMBA-treated *Rad51c^dah/dah^* **F-H)** Violin plots of MAPK signaling pathway genes (**F**) cytokine-cytokine receptor interaction pathway genes (**G**) and PI3K-Akt pathway genes (**H**) for male (blue) compared to female (pink) *Rad51c^dah/dah^* mice. **G)** RNAseq data heatmap of inflammatory pathways from KEGG analysis in (**D**) comparing DMBA-treated with non-DMBA-treated (CTRL) male *Rad51c^dah/dah^*.

Taking advantage of this strong gender-specific phenotypic difference seen with the *Rad51c^dah/dah^* mutations, we further analyzed the biological pathways induced by the transcriptional changes via KEGG enrichment analysis for 198 upregulated genes in male *Rad51c^dah/dah^* DMBA-treated group compared to females (Figs. 2C-2H, **Table S3**). This analysis revealed a gene expression program with higher expression of transcripts that belong to inflammation-related processes including Cytokine-cytokine receptor interaction (p=0.04) and IL-17 signaling pathway (p=0.04). Moreover, the MAPK signaling pathway (p=0.01), as well as PI3K-Akt signaling pathway (p=0.08) are upregulated, which are pathways associated with Collectively, the RNA-seq data analysis is consistent with strong inflammatory pathway upregulation and related macrophage activation.

### Western-like diet induces anemia of inflammation

Both inflammation and macrophage activation are characteristics associated with anemia of inflammation (AI) ^27,28^. Given our RNAseq results, we next sought to test if the observed anemia was AI. Macrophages have a prominent role in iron metabolism, as splenic macrophages remove excess iron from damaged and phagocytosed RBCs to recycle iron necessary for new RBC generation ^28,49^. Specifically, hepcidin as an iron metabolism hormone regulates iron by inhibiting the cellular export of Fe2+ which in turn leads to storage of the bio-inactive Fe3+ within the hemosiderin protein complexes. Hepcidin is transcriptionally controlled by IL-6, IL-1, IL-22, TGFb, IFNg, and TNFa. During AI, inflamed splenic macrophages are reprogrammed into granular macrophages, a distinct pathological hallmark of AI, similar to that seen in aging tissues. Hemosiderin complexes are crystalline structures that appear as distinct brown/grey pigments in hematoxylin and eosine (H&E) stained tissue sections, allowing the pathological identification of granular macrophages.

We next performed histological examination of spleens after chronic exposure to DMBA. H&E staining shows a high number of granular macrophages in the white pulp of male *Rad51c^dah/dah^*spleens, which is significantly greater than those found in female *Rad51c^dah/dah^*spleens (p=0.0013, Fig. 3A-3C). Reflecting the pattern of the RBC counts, spleens of male WT mice show the second largest number in granular macrophages between the groups, which is significantly larger than those seen in female WT spleen tissue (p=0.031, Fig. 3A-3C).

**Fig. 3.**
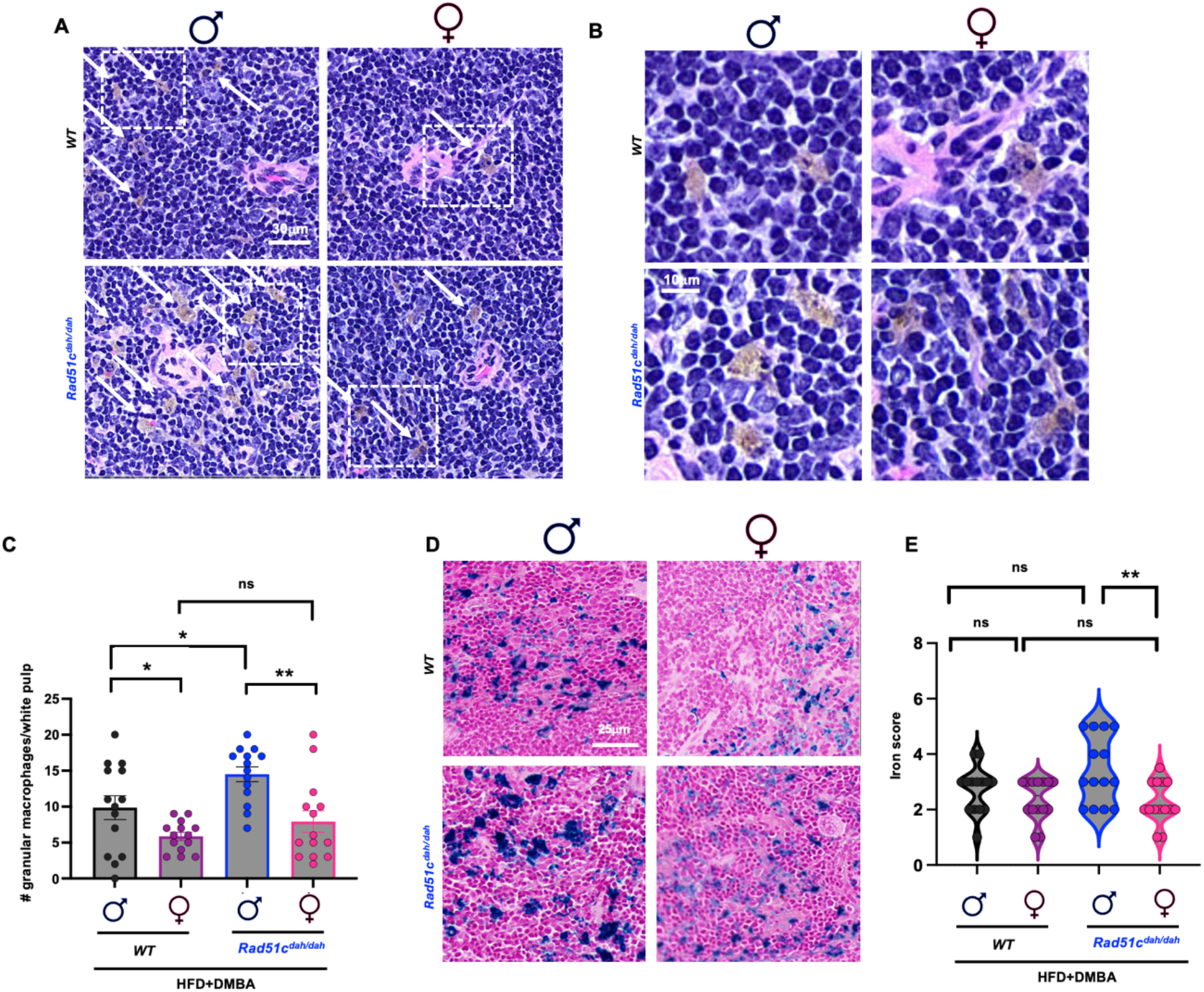
Male mice are prone to diet-induced anemia of inflammation. **A)** H&E stains of mouse spleen tissue sections (white pulp) of male (left) and female (right) WT (top) and mutant *Rad51c^dah/dah^* (bottom) mice treated with HFD+DMBA. White dashed boxes denote zoomed areas in (**B**), white arrows are pointing to granular macrophages **B)** Scatter bar plot of granular macrophage count in spleen (white pulp). Error bars denote standard deviation. n=14 per condition. **C)** Ferric iron stains of mouse spleen tissue sections of male (left) and female (right) WT (top) and mutant *Rad51c^dah/dah^*(bottom) mice treated with HFD+DMBA. **D)** Violin plot of (**D**) n=14 image fields for each condition. p-values are derived using the Student T-test. ns p>0.05, * p<0.05, **p < 0.01, ***p < 0.001, ****p<0.0001,

Hemosiderin triggers the release of iron from macrophages resulting in abnormal iron deposits in the surrounding tissue, another hallmark of AI pathology. Iron tissue deposits become unavailable for erythropoiesis, eventually resulting in anemia. To test for iron content directly we performed Perl’s blue staining in the spleens of male and female WT and mutant mice. The quantitation of iron deposits mirrors that of the granular macrophages, with male *Rad51c^dah/dah^* mice showing the highest amount of iron stain when treated with DMBA (p=0.0049 between male compared to female *Rad51c^dah/dah^*, Fig. 3D and 3E and Fig. S3A and S3B).

Inflammation driven macrophage reprogramming and bio-inactive iron overload are consistent with an AI phenotype.

### Male cells are vulnerable to mitochondrial DNA replication instability

DNA replication instability is a major source of inflammation ^15–18^. To investigate the observed sex-specific phenotype on a molecular level, we therefore examined DNA fork protection, a prominent DNA replication stability pathway that is important for suppression of inflammation and FA disease phenotypes ^13,14,45,50,51^. Specifically, the nascent DNA at damage-induced stalled replication forks requires protection from nucleolytic DNA degradation that otherwise triggers cGAS activation ^15,16^. Using single-molecule DNA fiber analysis, we measured the shortening of the nascent DNA label that is incorporated prior to replication stalling as a measure of fork degradation (Fig. 4A, ^13^). We have previously shown that when agnostic to the sex of the mice, mouse adult fibroblasts (MAFs) isolated from *Rad51c^dah/dah^* mice show a nuclear fork protection defect when challenged with hydroxyurea (HU), a nucleotide depleting agents that stalls DNA forks ^45^. To test potential gender-specific differences *in vitro* (Fig. 4) we isolated MAFs from both male and female mice. Consistent with previous results, *Rad51c^dah/dah^*mutant cells showed a nascent DNA tract shortening with HU in both male and female mutant MAFs (p<0.0001, respectively, Fig. 4B, blue and pink spheres). In contrast, WT male cells remained protected (p=0.0904, Fig. 4B, black spheres). Interestingly female WT cells exhibited a small but noticeable defect in fork protection, (0.8μm average change in tract length with and without HU in female WT cells, compared to 0.0μm in male WT cells). This trend was increased in mutant *Rad51c^dah/dah^* cells (4.1μm average change in tract length with and without HU in female WT cells, compared to 2.5μm in male WT cells), suggesting that female cells exhibit a greater nuclear replication instability compared to male cells. Overall, this suggests that genetics have the largest impact on nuclear fork instability under the conditions tested, and that nuclear instability does not explain the male sex bias seen with the *in vivo* phenotypes.

**Fig.. 4.**
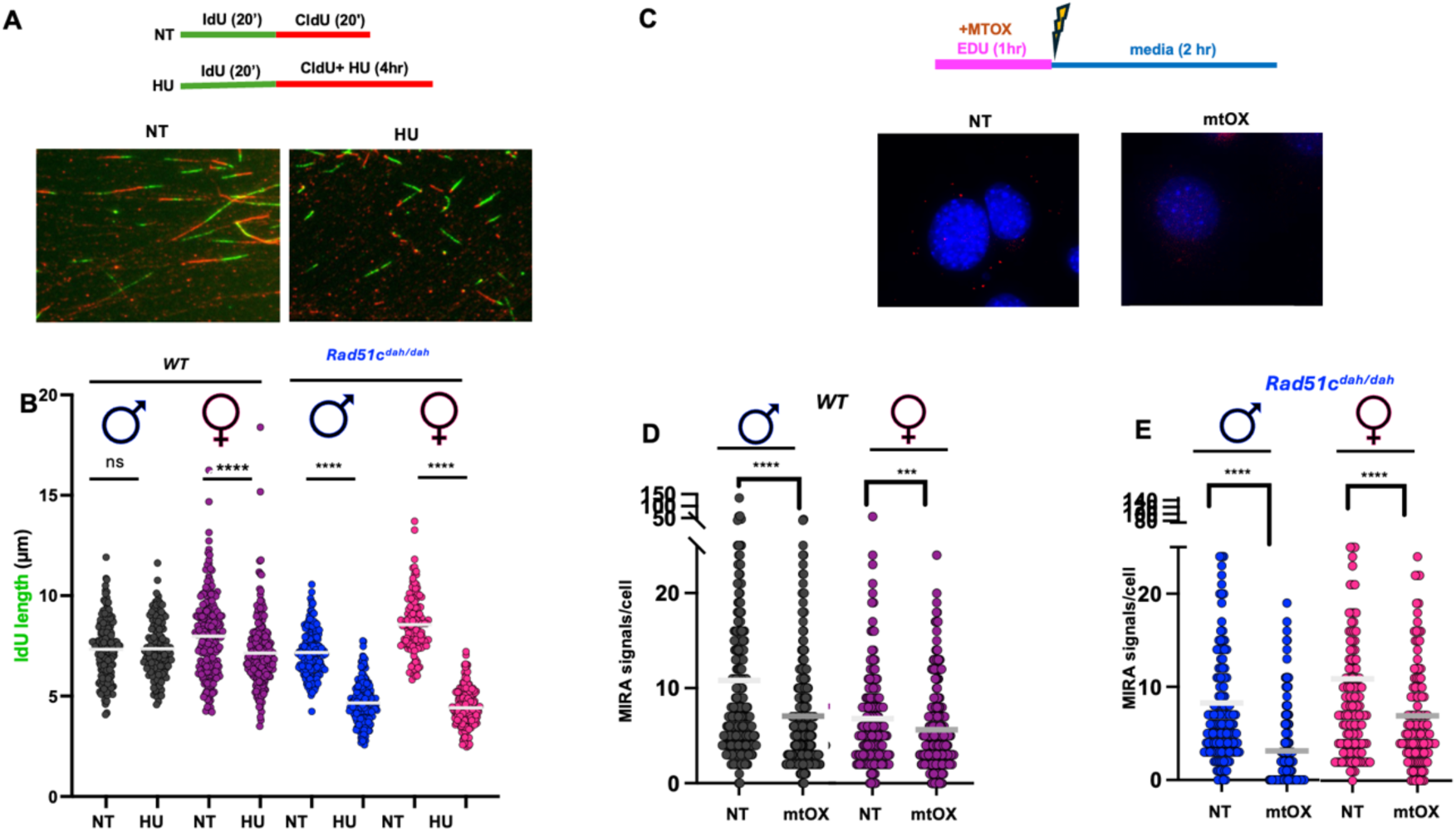
Male primary cells exhibit mitochondrial replication instability. **A)** Experimental sketch (top) and representative images (bottom) of single molecule DNA fiber spreads in primary mouse adult fibroblasts (MAF) with hydroxyurea (HU, 4mM, right image) and without (NT, no treatment, left image). Fork protection is measured as the loss of IdU tract signal (green) that had been incorporated before replication stalling with HU. **B)** Scatter plot of DNA fiber analysis of nascent IdU tract lengths, measuring DNA fork protection (n=125-222 from combined biological repeats). **C)** Experimental sketch (top) and representative images (bottom) of MIRA (mitochondrial replication assay) with EdU label incorporated immediately before oxidative damage by activation of MtOX (10μM), or no treatment (NT) in MAF cells **D-E)** Scatter plot of MIRA signals outside of nucleus in WT (**D**) and *Rad51c^dah/dah^* (**E**) MAF cells. p-values are derived using two-tailed Mann–Whitney test for DNA fiber analysis, and the Student t-test for MIRA analysis. ns p>0.05, * p<0.05, **p < 0.01, ***p < 0.001, ****p<0.0001, Bar represents the mean.

Both HFD and DMBA cause oxidative damage that can damage both nuclear DNA and mitochondrial (mt) DNA. BRCA/FA genes also play an active role in mitochondria to protect damaged mitochondrial DNA forks, which also suppresses a cGAS response ^20,25,26^. To measure mitochondrial fork protection, we employed a single-cell assay that measures the fate of mitochondrial DNA fork reaction (MIRA, ^20,52^). In MIRA, nascent mitochondrial DNA is marked using a nucleotide analogue incorporated into newly synthesized mtDNA, and a proximity ligation assay is used for signal amplification of the nascent DNA. Upon treatment with mtOX, an oxidative damaging agent specifically targeted to the mitochondria to avoid cross-communication with nuclear damage ^20,53^, MIRA signals are markedly reduced in male mutant *Rad51c^dah/dah^* cells, indicating a loss of nascent mtDNA and mitochondrial fork instability (p<0.0001, Fig. 4C-4E, and Fig. S4). While female mutant *Rad51c^dah/dah^* cells do lose some of the mtDNA signals under these treatment conditions, this loss is considerably less pronounced (Fig. 4E). The trend was similar in WT cells, with male WT cells showing more pronounced mitochondrial fork instability compared to female WT cells (Fig. 4D). Collectively, the data reveals that in contrast to the nucleus, male cells are more susceptible to mitochondrial DNA replication instability.

### The Un-P-STAT1 response is preferentially induced in male mice

DNA replication instability produces DNA fragments that stimulate the cGAS-STING Type I interferon (IFN) pathway. While nuclear instability resulting in micronuclei mediated cGAS activation have been shown to promote interferon production as part of the canonical Type I IFN response leading to JAK/NFKb pathway activation ^15–18^, we and others recently showed that mitochondrial DNA instability promotes a different Type I IFN response ^20,25,26^. Faulty mitochondrial fork protection, mitochondrial transcription factor A (TFAM) or mutant DNA polymerase gamma (POLG) mediated mitochondrial instability all activate the expression of a distinct subset of cGAS-dependent interferon stimulated genes (ISGs), which constitute the Un-P-STAT1 response (also termed ISG3 response) ^20,25,26^. Unlike the canonical Type I IFN, the un-P-STAT1 response does not induce IFNa/b or JAK/NFKb pathway targets ^22,23^. Given that mtDNA fork protection suppresses this distinct pathway and showed a sex bias, we sought to test if the mtDNA-dependent Un-P-STAT1 response similarly is controlled in a sex-specific manner.

Further analyzing our RNAseq data for the Un-P-STAT1- gene expression signatures ^20,26^, we found substantial enrichment of ISG gene transcripts in the male *Rad51c^dah/dah^* DMBA- treated compared to untreated mice (Fig. 5A). Similarly, the Un-P-STAT1 signature is enriched when comparing male to female Rad51c^dah/dah^ mice (Fig. 5B). Consistent with a primary Un-P- STAT1 response, JAK pathway transcripts are not enriched, but rather reduced with DMBA treatment and in males compared to females (Fig. S5A and S5B). Employing this refined inflammation pathway signature analysis revealed a notable difference in expression of transcripts of the Un-P-STAT1 pathway also in wild-type mice, with males showing enrichment in this gene signature gene compared to females (Fig. 5C) as well as in males comparing control with DMBA treatments These data suggest that the Un-P-STAT1 response is linked to the AI phenotype observed in both wild-type and mutant male mice.

**Fig. 5.**
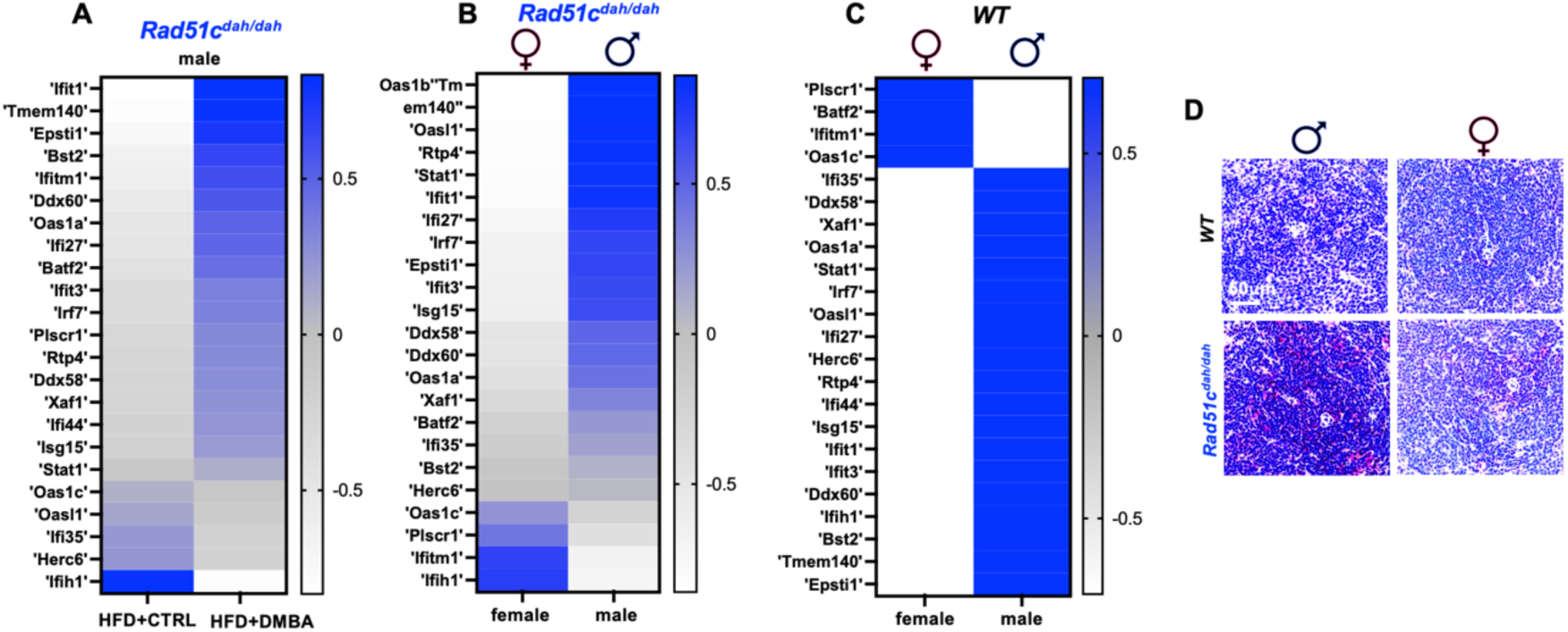
Activation of the un-phosphorylated STAT1 response is stronger in male mice A-C) RNAseq data heatmap of un-phosphorylated STAT1 response pathway in in bone marrow of HFD fed mice comparing (**A**) male *Rad51c^dah/dah^* mice with and without DMBA (**B**) male with female *Rad51c^dah/dah^* mice with DMBA and (**C**) male with female *WT* mice with DMBA. **D)** Representative images Red Alkaline phosphatase stains against STAT1 in mouse spleen tissue sections (white pulp) of male (left) and female (right) WT (top) and mutant *Rad51c^dah/dah^* (bottom) mice treated with HFD+DMBA. Scale bar, 50μm

In contrast to the canonical cGAS/JAK Type I IFNa/b response activated by phosphorylation of STAT1 at Tyrosine 701 to promote the inflammation signaling cascade, the mt-DNA dependent Un-P STAT1 response shows negligible changes in P-Tyr701 STAT1 levels. Instead, total STAT1 protein levels are increased in the Un-P-STAT1 response as a distinguishing feature ^20,22,23,26^, whereas total STAT1 typically remains unchanged during the canonical Type I IFN response. To directly test if the Un-P-STAT1 response is stimulated by the diet regimens in a sex-specific manner, we assessed total STAT1 protein in murine spleens (Fig. 5D, and Fig. S5D and S5E) using immunohistochemistry. In the male DMBA treated *Rad51c^dah/dah^* mice, STAT1 protein levels are enhanced compared to female *Rad51c^dah/dah^* mice (Fig. 5D), as well as when compared to DMBA-untreated mice (Fig. S5E). These data show that the mtDNA dependent Un-P-STAT1 response is induced under chronic stress conditions that promote anemia of inflammation.

### Estrogen protects against stress-induced mitochondrial DNA replication instability

To further dissect the molecular mechanism by which females may be protected against induction of the Un-P-STAT1 response, we sought to better mimic the diet induced stresses *in vitro* and treated male and female mutant and WT MAFs with DMBA (Fig. 6A-6C). DMBA is known as a DNA intercalating agent. In mitochondria, DMBA generates reactive oxygen species (ROS, ^54^), which is known to cause lesions that potently stall mitochondrial DNA replication ^55^. Using the MIRA assay, the data shows a stark loss in nascent MIRA signals in both WT and mutant male cells (Fig. 6A-6C). In contrast, mitochondrial fork degradation in female cells is much less pronounced (Fig. 6A-6C), suggesting protection against DMBA induced loss of mtDNA in female cells. Treatment of cells with DMBA and visomitin, a mitochondrial targeted antioxidant ^56,57^ rescues the MIRA signal loss in both male WT and male mutant cells (Fig. 5A- 5C), suggesting that induction of ROS is the cause for MIRA signal loss with DMBA. In stark contrast, MIRA signals remained unchanged in female cells.

**Fig. 6.**
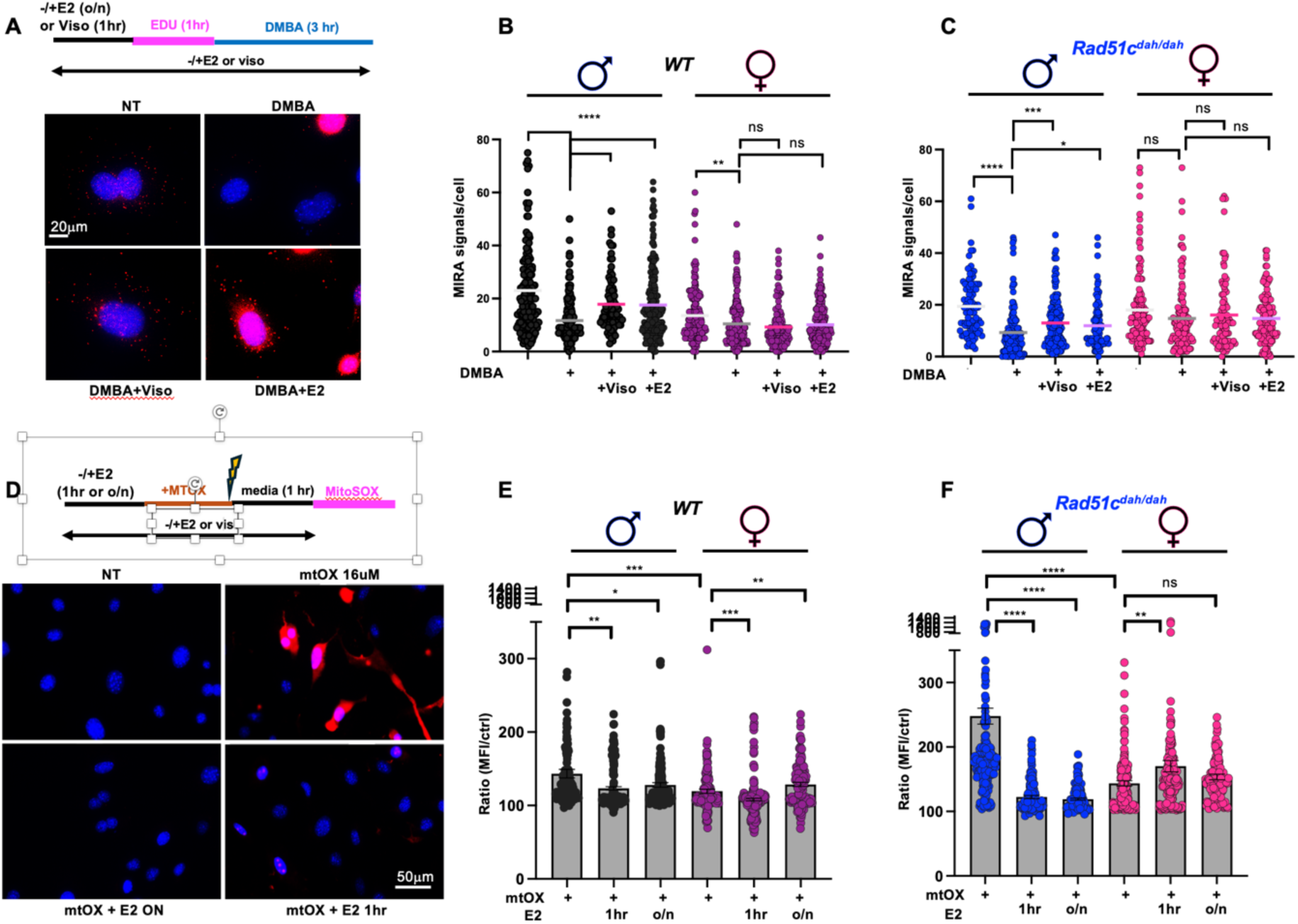
Estrogen protects from mitochondrial replication instability and damage. **A)** Experimental sketch and representative images of MIRA in male *Rad51c^dah/dah^* MAF cells. EdU label is incorporated immediately before treatment with DMBA (6μM), with or without Visomitin (viso, 500nM) or Estrogen (E2, 10μM), or no treatment. **B-C)** Scatter plot of MIRA signals outside of the nucleus in WT (**B,** n=88-220) and *Rad51c^dah/dah^* (**C,** n=46-156) MAF cells. Bars represent the mean. **D)** Representative images of mitochondrial superoxide staining in male *Rad51c^dah/dah^* MAF cells using MitoSOX with or without Estrogen (10μM) for one hour prior or 24 hours (ON, overnight) prior to exposure to MTOX (16μM) as indicated in the experimental sketch. E2, estrogen; mtOX, light-activated mitochondrial targeted oxidative damage; ON, overnigh **E-F)** Scatter plot with bar of MitoSOX signal intensity in WT (**E,** n=152-324) and *Rad51c^dah/dah^* (**F,** n=139-244) MAF cells. Bars represent the mean with SEM.

One of the most prominent molecular differences between male and female organisms and cells is the enrichment of estrogen in females. As estrogen receptors are present on both nuclear and mitochondrial plasma membranes ^58,59^, we next sought to define the molecular effect of estrogen on mitochondrial fork instability. Strikingly, treatment of male cells with estrogen was observed to rescue the effect of DMBA in male WT and mutant *Rad51c^dah/dah^* cells, and significantly increase MIRA signals compared to DMBA treatments alone in both WT and mutant *Rad51c^dah/dah^* cells. Thus, estrogen protects against mitochondrial fork degradation, providing a molecular explanation for the suppression of the Un-P-STAT1 response seen in females.

While estrogen can activate a signature transcriptional program by binding membrane bound estrogen receptor, estrogen has also been reported to act as an antioxidant via both transcription-dependent and transcription-independent mechanisms ^58,60,61^. We therefore sought to measure the impact of estrogen on mitochondrial ROS in our cells (Fig. 6D-6F) by measuring the extent of ROS using mitochondrial superoxide indicator (mitoSOX), a fluorogenic dye specifically targeted to mitochondria in live cells (Fig. 6D-6F). While marginal mitoSOX signal was detected in untreated cells, the addition of mtOX significantly induced ROS in both mutant and WT cells (Fig. 6D-6F and Fig. S6). Comparing the male and female cells, the same mtOX treatment induced significantly more mitochondrial ROS in both male WT and mutant cells compared to female cells (p=0.0005 and p<0.0001, respectively, Fig. 6D-6F) suggesting female cells are protected against mitochondrial oxidative damage. Adding estrogen prior to mtOX significantly lowered mitoSOX intensities in WT and *Rad51c^dah/dah^* male cells (p=0.002 and p<0.0001, respectively, Fig. 6D-6F). Of note, estrogen lowered mitoSOX signals in male cells irrespective of whether estrogen was added the day before the experiment or just one hour prior to mtOX treatment, which is a time-frame that has been shown to only have negligible effects on estrogen-induced transcriptional changes compared to long-term treatments ^62,63^. In sum, the data suggest that estrogen can act as a mitochondrial antioxidant, resulting in male cells being more vulnerable to mitochondrial ROS and associated mitochondrial DNA fork instability.

### Mitochondrial fork protection defects contribute to anemia of inflammation

Our data demonstrates that male WT, and to a greater degree male *Rad51c^dah/dah^*mice, develop anemia under chronic stress conditions, a phenomenon that is associated with the mitochondrial DNA stress-induced Un-P-STAT1 inflammatory signaling response. On a cellular level, males have higher mitochondrial oxidative stress and consequentially greater mitochondrial replication fork instability that is preventable by exogenous estrogen. These observations suggest that the anemia in the male mice is preventable by various selective treatment agents, including estrogen, agents reducing inflammation processes, and by oxidative damage inhibitors specifically targeting the mitochondria.

To test if the molecular insights from the cellular data are translatable *in vivo*, we compared chronically HFD+DMBA treated male *Rad51c^dah/dah^*mice with those treated with estrogen before and during the DMBA treatment regime (Fig. 7A). Strikingly, complete blood analysis reveals that estrogen is sufficient to protect against HFD+DMBA-induced anemia (p=0.0028, Fig. 7B-D and Figs. S7A-E). Interestingly, the protective effect of estrogen is restricted to red blood cells, whereby white blood cells and platelets are markedly albeit insignificantly reduced with the additional estrogen treatment (Fig. 7E and 7F).

**Fig. 7.**
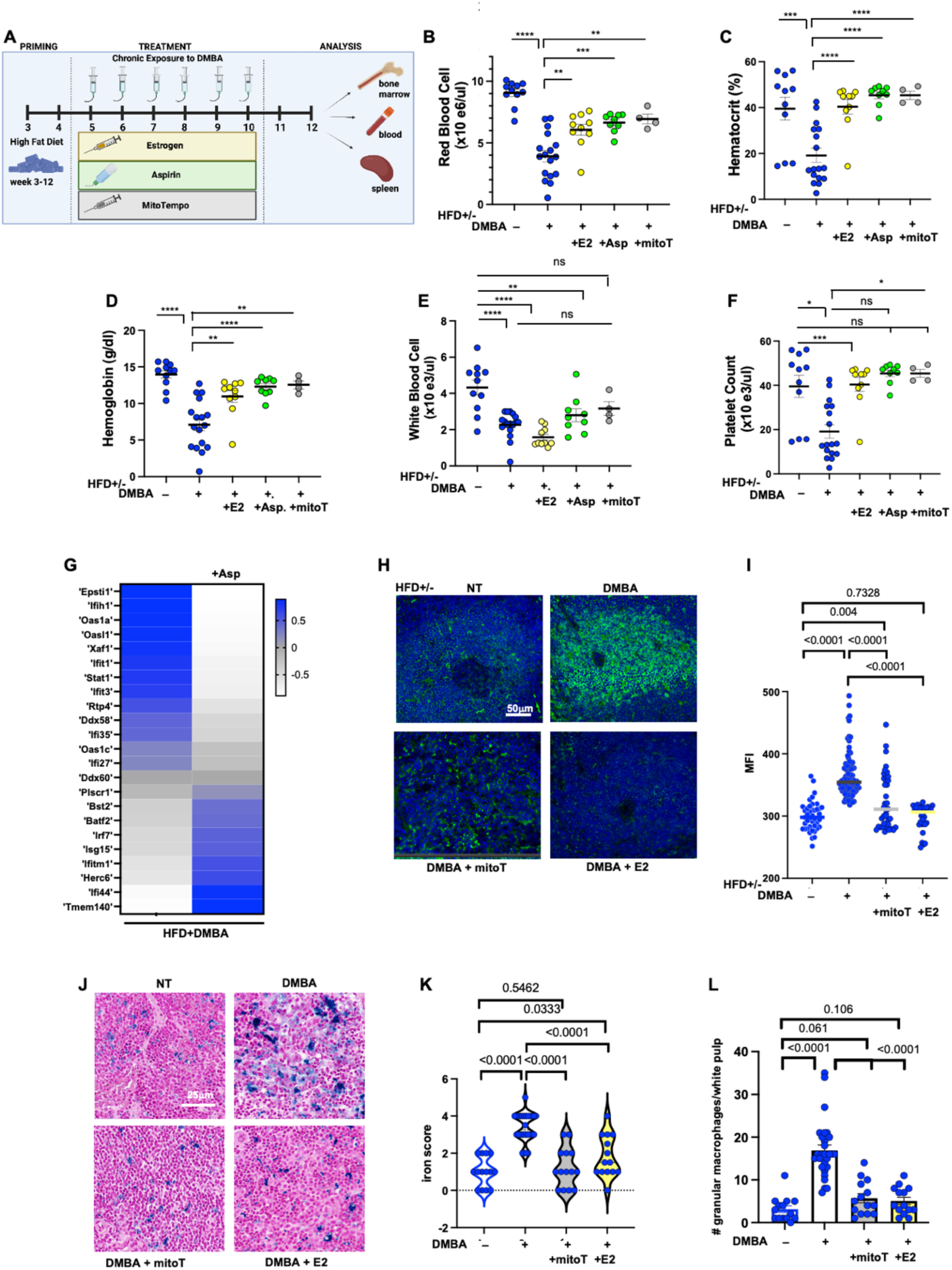
Anemia in male *Rad51c^dah/dah^* under chronic stress exposure is suppressed by restoring mitochondrial stability. **A)** Experimental workflow for rescue mouse experiments under chronic stress treatment with estrogen (yellow), aspirin (green) and mitoTempo (grey) **B-F)** Red cell blood count (**B**) hematocrit (**C**), hemoglobin (**D**), white blood cell (**E**), and platelet count (F) from male *Rad51c^dah/dah^* mice with or without DMBA-treatment (blue) and rescued with additional treatment with estrogen (yellow), aspirin (green) or mitoTempo (grey). **G)** Heatmap RNAseq data heatmap of un-phosphorylated STAT1 response pathway in bone marrow of HFD fed mice comparing male *Rad51c^dah/dah^* mice with DMBA with and without additional treatment with aspirin. **H)** Representative images immunofluorescence stains against STAT1 in male *Rad51c^dah/dah^* mouse spleen tissue sections (white pulp) treated with HFD+DMBA, with or without additional treatment of mitoTempo or estrogen (E2). Scale bar, 50mm **I)** Scatter plot of quantitation of **H** (n=21-89). MFI, mean fluorescence intensity; bar denotes median. **J)** Ferric iron stains of male *Rad51c^dah/dah^* spleen tissue sections treated with HFD+DMBA with or without additional treatment of mitoTempo or estrogen (E2) **K)** Violin plot of (**J**) n=14-28 image fields for each condition. **L)** Scatter bar plot of granular macrophage count in spleen (white pulp). Error bars denotes the SEM. n=14-28 per condition. p-values are derived using the two-tailed Student T-test. ns p>0.05, * p<0.05, **p < 0.01, ***p < 0.001, ****p<0.0001,

Aspirin is a non-steroidal anti-inflammatory drug (NSAID) widely used to relieve pain and inflammation, which more recently has been explored in diseases associated with chronic inflammation such as cancer, albeit its efficacy may be context dependent ^64^. To validate the involvement of chronic inflammation processes driving the anemia, we fed our animals continuously with low concentration of aspirin in their drinking water (Fig. 7B-F and Figs. S7A-E). Treatment with aspirin rescues anemic mice (p=0.0002) and improves WBC and platelet counts, suggesting that aspirin may target the Un-P-STAT1 response. Consistent with the latter, aspirin reverses the induction of the Un-P-STAT1 response at the RNA level (Fig. 7G and Figs. S7F-J).

To further confirm that mitochondrial instability contributes to inflammation-induced anemia, we used a mitochondria-specific ROS scavenger to prevent the mitochondria damage by DMBA-induced ROS (Fig. 7A). Treatment of male *Rad51c^dah/dah^*with mito-TEMPO, an antioxidant specifically targeted to the mitochondria ^65^ rescues RBC and platelet count (p=0.0026 and p=0.0171, respectively), and improves WBC count (Fig. 7B-F and Figs. S7A-E). Of note, some of the mitoTEMPO+HFD+DMBA but none of the mitoTEMPO+HFD treated animals developed skin lesions, which were excluded from the blood analysis as the DMBA- induced lesions are likely cancerous, which would skew the blood count results. Nevertheless, spleen tissue sections in non-cancerous mitoTEMPO treated mice shows a reduction in total STAT1 protein, iron deposits and granular macrophages, similar to estrogen treatment (Fig. 7H-7L), suggesting that mitochondrial ROS drives the anemia.

Taken together the data show that mitochondrial DNA replication instability inducible by unfortunate diet causes a chronic inflammation response driven by Un-p-STAT1 signaling that contributes to anemia of inflammation.

## Discussion

### Sex specific differences in inflammation-driven disease

The importance of gender differentially driving disease etiology has recently emerged as a critical determining factor. Anemia of inflammation (AI) exemplifies this, with aged men and male diabetic patients exhibiting AI markedly more frequently compared to women ^27–29^. The underlying molecular mechanisms for these differences have proven to be elusive. Our data identifies mitochondrial replication instability as a key molecular driver for gender specific differences in inflammatory responses driving AI, a phenomenon that is particularly pronounced in the context of existing genetic predisposition, as seen in the *Fanco/ Rad51c^dah/dah^* mouse model.

Specifically, our data suggest that estrogen exerts its protective effect by reducing oxidative stress in the mitochondria. Mitochondrial oxidative stress in turn drives mitochondrial replication instability that activates the Un-P-STAT1 response, which drives the anemia. Estrogen has been reported to upregulate the transcription of antioxidant pathways ^66,67^. Based on the short timeframe needed for estrogen to take effect in our disease model, our data support that estrogen might exert its antioxidant function in part through non-transcriptional pathways. Our conclusions extend previous reports in the literature ^58,60,68^.

The mechanistic basis of estrogen’s function as a molecular antioxidant has yet to be defined, and has been proposed to involve estrogen’s phenol properties or MnSOD activation ^58,60,68^. Our data now suggest a novel aspect, implying that estrogen has mitochondria-specific antioxidant functions that promote mitochondrial genome stability. Biologically, this could be a reflection of the matrilineal inheritance of mitochondria. Intriguingly, our data show that while on the one hand estrogen promotes mitochondrial DNA replication stability and suppresses associated inflammation reactions, there is a noticeable decrease in nuclear fork protection in female cells compared to male cells under identical experimental conditions. The observed estrogen-dependent nuclear genome instability is consistent with previous reports that show estrogen promotes transcription-replication collisions ^69^.

Further support of this observation is provided by our *in vivo* results; while estrogen treatment in mice rescues the anemia in the male *Rad51c^dah/dah^*mice, it exacerbates the loss of white blood cells and platelets. The latter two cell types contain a nucleus while red blood cells do not. The nuclear fork protection defect may cause depletion of cells that have a nuclear genome under these conditions, which would explain the differential response of the three blood lineages to estrogen treatment.

Sex bias in disease is not restricted to anemia. For instance, males are more susceptible to COVID infection ^70^ and cardiac disease ^71^. Overall males exhibit a shorter lifespan irrespective of ethnicity or demographics in most parts of the world, with the main contributing factor suggested to be cardiac disease ^72^. Intriguingly, the mitochondria-DNA dependent Un-P-STAT1/ISG3 response has been linked to cardiac toxicity and cardiac disease ^24^ and is involved in the antiviral response ^26^. Thus, it is plausible that the increased capacity of female mitochondria to suppress this distinct Type-I interferon signaling pathway may also contribute to improved suppression in diverse diseases.

### AI is driven by accumulated stress

The AI phenotype and the difference between males and females is more pronounced in the *Fanco*/*Rad51c^dah/dah^*mice compared to wild-type mice. It is important to note that the *Rad51c^dah/dah^*mouse model does not develop FA, anemia or any other noticeable phenotypes unless specifically challenged. Thus, it exemplifies how additional genetic components can influence and skew disease outcome. In principle, it is possible that some of the results are mutant specific. Alternatively, it is a reflection of phenotypic presentation with increased amounts of stress, which can be caused by external or internal damage, or by stress induced by genetic defects. In support of a stress-dose dependence, our data shows that DMBA treatment alone in the absence of a HFD is insufficient to significantly promote anemia in wild-type males, while the mutant male mice exhibit mild anemia, together suggesting that accumulation of stress gradually worsens the outcome. In this context, we have previously shown that a *FANC* gene mutation causes sufficient genetic stress to exacerbate replication stress responses and consequently development of disease ^45^. Importantly, an estimated ∼78% of the Western population harbor at least one disease-associated heterozygous FANC variant ^44^. Since heterozygous mutations in FANC are sufficient to cause fork protection defects ^73,74^, the results reported here likely are relevant for a majority of the population.

### The emerging importance of the UN-P-STAT1 response

Our data highlight the importance of the Un-P-STAT1 response, which is induced by chronic, low grade stress ^21–23,25,26^. Recent studies have implicated the Type I IFN sub-pathway in antiviral defense, inflammation-induced chemo-therapy resistance in cancer cells ^25,26^, chemotherapy induced cardiac toxicity and cardiac disease ^24^, highlighting both the universality and tissue context specificity of the response. Our results extend the current understanding of the Un-P-STAT1 response and reveal that it contributes to macrophage reprogramming and iron metabolism rewiring, resulting in AI. Importantly, females are protected against the Un-P-STAT1 response, distinctively expanding our understanding of differences in sex-dependent inflammation response pathways and how these are achieved on a molecular level.

Multiple cytokine signaling pathways are known to be important in the development of AI, including the inflammasome-dependent IL1, Toll-like receptor pathway IL6, TGFb pathways, TNFa and Type II IFNg pathways. Intriguingly, TGF-b also has been implicated in promoting pancytopenia by HSC exhaustion ^75^. AI is not reported to be associated with pancytopenia. Nevertheless, one or more of the other blood cell compartments are affected in multiple chronic disease and stress situations that promote AI including Crohn’s disease, diabetes, aging, and cancer therapy response. Our data show that AI caused by diet induced stress is accompanied by mild pancytopenia. These results suggest that considering blood cell compartments other than red blood cells may be an informative additional parameter in assessing the severity of AI.

Aspirin, and restoration of mitochondrial fork protection with mitochondrial-targeted antioxidants or estrogen suppresses the Un-P-STAT1 response, reverse AI and macrophage reprogramming in our animal model, magnifying the importance of this emerging inflammatory signaling pathway. It is currently not clear if other AI associated cytokine pathways are a consequence of the Un-P-STAT1, a cause or an independent event. Direct communication between cytokine pathways has been reported including a cGAS dependent inflammasome responses ^76^. It will be of great importance to dissect these relationships in future studies to understand if mitochondrial fork instability can independently of the Un-P-STAT1 response activate diverse pathogens or DNA damage signal stimulated pathways including inflammasome, toll-like receptor and Type II IFN response pathways. The data further show that adverse diet induces sufficient stress to dysregulate the Un-P-STAT1 response. Future studies are needed to test if a beneficial diet could suppress or normalize it.

The Un-P-STAT1 response so far has been consistently associated with mitochondrial DNA stress ^24–26^. In principle, it is possible that it also can be triggered by nuclear DNA stress under certain circumstances. However, given its small size, we suggest that the mitochondrial genome is more vulnerable to low amounts of damage compared to the nuclear genome, positioning mitochondria as an ideal first responder in the cellular communication of low-dose, chronic stress responses.

Collectively, the results here call for an increased awareness of organelle-specific genomic instabilities that can drive sex-specific inflammation and disease.

## ACKNOWLEDGEMENTS

We thank Drs. John Tainer and Davide Moiani for discussions on RAD51C structures. We thank the technical support from the Cancer Prevention and Research Institute of Texas (CPRIT RP180734) and Dr. Hariyadarshi Pannu for feedback on the manuscript. This work was supported by the Cancer Prevention and Research Institute of Texas and NIEHS under award 1R01ES029680, and by CPRIT RP180463, R1312 and RP180813 (K.S.). K.S. is a Rita Allen Foundation Fellow and a CPRIT scholar in Cancer Biology (previous award R1312).

## AUTHOR CONTRIBUTIONS

*Conceptualization,* K.H.T. and K.S*.; methodology and investigation:* K.H.T., Y.C., S. R., M. H., C.L.T. *J. Z., writing* K.H.T. and K.S*.; funding acquisition,* K.S.

All authors discussed the results, read and approved the manuscript.

## DECLARATION OF INTERESTS

The authors declare no competing interests.

p-values are derived using the two-tailed student T-test for MIRA analysis. ns p>0.05, * p<0.05, **p < 0.01, ***p < 0.001, ****p<0.0001

## Supplemental information

Document S1. Figs. S1-S7 and Table S1-S4

## METHODS

### Institutional review board statement

All procedures and methods were conducted in accordance with federal and state regulations as well as MD Anderson Cancer Center institutional guidelines and policies. All *procedures performed* on *animals* was described in an *Animal Care* and Use Form (ACUF) and approved by the *institutional animal care and use committee* (IACUC).

### Treatment of mice

Generation of *Rad51c^dah/dah^* mice on a pure FVB/J background was previously reported ^45^. Mice were maintained in pathogen-free conditions. Animal experiments were performed per approved animal protocol by the Institutional Animal Care and Use Committee of the University of Texas M. D. Anderson Cancer Center. Three-week-old mice were fed with free access to rodent chow (10% of total calories from fat, control diet) and water or high fat diet (HFD, Table S1) (60% of total calories from fat) until endpoints. The mice were separated into groups as follows: control diet (con, *n* = 6-7 per group), control diet with 7,12-Dimethylbenz(a)anthracene (DMBA) treatment (con+DMBA, *n* = 6-7 per group), HFD group (HFD, *n* = 6-8 per group); and HFD + DMBA (*n* = 6-11 per group), HFD+ estradiol treatment (*n* = 7 per group); and HFD+DMBA+estradiol treatment (*n* = 10 per group), HFD+ aspirin treatment (*n* = 7 per group); and HFD+DMBA+aspirin treatment (*n* = 9 per group), HFD+mito-TEMPO treatment (*n* = 6 per group); and HFD+ DMBA+mito-TEMPO treatment (*n* = 4-7 per group), DMBA was administered by oral galvage (25 mg/kg, once a week for 6 weeks) starting at 5 weeks of age. Non-DMBA treated mice received corn oil. Estradiol was administered subcutaneously 4h before DMBA treatment (0,1 ug, once a week for 6 weeks) and provided via drinking bottle with a final concentration of 4ng/ml from week 4-12. Aspirin were provided via drinking bottle with a final concentration of 0,3mg/ml from week 4 onwards. Mito-TMEPO were administered intraperitoneal (2mg/kg, twice a week for 6 weeks) starting at 5 weeks of age. Body weights were measured over the treatment period. At 5, 6 or 12 weeks mice were euthanized and rapidly prepared for downstream analysis.

### Histological analysis

Spleens from treated mice were resected and fixed in 10% buffered formalin solution (Thermo Fisher Scientific) overnight and stored in 70% ethanol. Formalin-fixed tissues were paraffin-embedded and tissue sections were counterstain with Hematoxylin and eosin using standard procedure on the Gemini Auto stainer. Ferric iron staining was performed using kit KTIRO from StatLab, according to manufacturer’s instructions. AP/Red IHC staining was performed using DoubleStain IHC Kit from Abcam, using manufacturer recommendations. STAT1 (D1K9Y, cell signaling technology) antibody was used at a dilution of 1-400. Images were obtained using a Nikon Eclipse Ti-U inverted microscope with an Andor Zyla sCMOS camera and scanned using Pannoramic midi slide scanner (3DHistech). For iron stain scoring, image fields were analyzed using a binned approach against an internal reference scale (1-5 with 1 having the least and 5 the most amount of iron stain amongst the samples).

### Immunofluorescence

Formalin-fixed, paraffin-embedded (FFPE) sections of mouse spleen were stained by immunoflurescence using 1-400 dilution of anti-STAT1 (D1K9Y, Cell signaling technology). Briefly, slides were deparaffinized, rehydrated and antigen retrieval was performed by citrate buffer. Slides were blocked for 1 hour at room temperature and incubated with primary antibody overnight. Sides were washed in TBST buffer the following day and stained with secondary antibody conjugated to anti-rabbit Alexa fluor 488 (Thermo Fisher Scientific) for 1 hour, washed and counter-stained for DAPI prior to mounting with Prolong gold antifade (Thermo Fisher Scientific). Images were obtained using a Nikon Eclipse Ti-U with an Andor Zyla sCMOS camera and scanned using Pannoramic midi slide scanner (3DHistech). Images were analyzed using the Nikon NIS Elements. Statistical analysis was performed using GraphPad Prism 10 software.

### RNA-Sequencing and Sample Preparation

Femur and tibia were collected from experimental mice and bone marrow single-cell suspension were collected by flushing tibias and femurs with ice-cold PBS. Single-cell suspension were collected in RNALater and directly fresh-frozen in liquid nitrogen. RNA was isolated from bone marrow samples using RNeasy Micro Kit (Qiagen) according to the manufacturer’s instructions. Additionally, excessive mRNA from blood cells were removed using globulin gene removal GLOBINclear kit (Invitrogen) according to the manufacturer’s instructions. RNA samples of multiple mice per condition were submitted to Cancer Genomics Center at The University of Texas Health Science Center at Houston for sequencing. Total RNA was quality-checked using Agilent RNA 6000 Pico kit (#5067-1513) by Agilent Bioanalyzer 2100 (Agilent Technologies, Santa Clara, USA). RNA with an integrity number of greater than 7 was used for library preparation. Libraries were prepared with KAPA mRNA HyperPrep (KK8581, Roche) and KAPA UDI Adapter Kit 15μM (KK8727, Roche) following the manufacturer’s instructions. The quality of the final libraries was examined using Agilent High Sensitive DNA Kit (#5067-4626) by Agilent Bioanalyzer 2100 (Agilent Technologies, Santa Clara, USA), and the library concentrations were determined by qPCR using Collibri Library Quantification kit (#A38524500, Thermo Fisher Scientific). The libraries were pooled evenly and went for the paired-end 75-cycle sequencing with 25 million reads for each end per sample on an Illumina NextSeq 550 System (Illumina, Inc., USA) using High OutputKit v2.5 (#20024907, Illumina, Inc., USA). BGI Genomics Inc performed analysis by using DNBSEQ platform to filter reads (PE75 sequencing length), the HISAT platform to align the clean reads to the reference genome (mus musculus, GCF_000001635.26_GRCm38.p6), and Bowtie2 to align clean reads to reference genes (BGI Genomics Inc.). Heatmaps were created using GraphPad PRISM 10 comparing between treatment conditions by scaling by row ^77^ and averaging biological replicas, and pathway analysis was performed using the software tool Dr. Tom Data Visualization Solution (BGI Genomics Inc.).

### Peripheral blood analysis

Whole blood was collected from indicative treatments and appropriate controls in EDTA microvette tubes (BD Biosciences) and complete blood counts were analyzed using ADVIA 2120i Hematology systems (Siemens Healthineers) according to the manufacturer’s instruction. Whole blood slides were stained with Diff-Quick stain.

### Generation of mouse adult fibroblasts (MAF)

Primary ear fibroblasts were derived from age-matched males and females. Briefly, a portion of the ear was cut off, rinsed two times with PBS containing kanamycin (100 μg/mL) and digested with collagenase D/dispase II protease (4 mg/mL, respectively) for 45min at 37°C. After dilution with five times Dulbecco’s modified eagle’s high-glucose media (DMEM) containing 10% fetal bovine serum (FBS) and 5% Antibiotic-Antimycotic, cells were incubated overnight at 37°C. The following day, cells were passed through a 0.7μM cell strainer, washed with PBS and plated for cultivation in standard media consisting of DMEM supplemented with 10% FBS and 100units/ml Pen-Strep. MAFs were grown at 37°C and 5% CO2, routinely passaged two times per week and passages 2 to 10 were used for experiments.

### DNA Fiber Assay

DNA fiber assays were performed as previously described ^14^. Briefly, MAF cells were seeded in a 6-well plate at a density of 1×10^5^ cells per well. The next day, they were pulse-labelled with 50 μM 5-iodo-2’-deoxyuridine (IDU, Sigma #I7125) for 20 minutes. After IDU treatment, the cells were washed 2-3 times with warmed PBS followed by 50 μM 5-chloro-2’-deoxyuridine (CldU, Sigma #C6891) for 20 minutes or 50 μM CldU with 4mM hydroxyurea (HU) (Sigma #H8627) treatment for 4 hours. Cells were harvested by trypsinizing, resuspended in ice-cold PBS and spotted onto glass slides and lysed in 200 mM Tris-HCl, pH 7.4, 50 mM EDTA, 1% SDS. DNA was allowed to attach for 5.5 minutes before spreading by gravity. Slides were fixed in methanol/acetic acid (3:1). Slides were air-dried at room temperature overnight and denatured in 2.5 M HCl for 30 minutes followed by neutralization with PBS (pH 8, and subsequent pH 7.5 washes). Slides were blocked with 5% BSA and 0.1% Triton X in PBS. IdU/CldU fibers were stained using standard immunostaining with antibodies against IdU (anti-BrdU, clone B44, Beckton Dickinson #347580, 1:50) and CldU (anti-BrdU BU1/75, Novus Biological ab6326, 1:100) followed by sequential labelling with secondary antibodies (Invitrogen, goat anti-mouse AF488; 1:100 and goat anti-rat AF555, 1:200) before mounting slides with Prolong Gold (Invitrogen, P36934). IdU/CldU Fibers were imaged using a Nikon Eclipse Ti-U inverted microscope and analyzed using ImageJ software.

### In situ quantification of nascent mitochondria DNA assay (MIRA)

Mira assays were conducted as previously described ^20,52^. Briefly, MAF cells were plated in 8-well chamber microscope slides (Thermo Scientific, 177402) one day before treatment. The next day, media was exchanged with DMEM containing 10% charcoal stripped FBS (Gibco, 12676-029) and incubated overnight with or without 10 μM Estrogen (Sigma, E8875). The cells were treated with 20 μM 5-ethynyl-2′-deoxyuridine (EdU, Invitrogen, A10044) for 1 hour in the presence or absence of 10 μM mtOX ^53^ or 6 μM DMBA (Selleck Chemicals, E1022), 500 nM Visomitin (Medchemexpress, HY-100474) or 10 μM Estrogen for the indicated time, followed by incubation in ambient light for 5 minutes to allow for mtOX activation. Cells were washed in PBS and allowed to recover in medium as indicated in the Figs.. Cells were fixed with 4% paraformaldehyde (EMS, 15714), permeabilized with 0.25% Triton X-100/PBS and click-iT reaction was performed using biotin azide (Invitrogen, B10184) and AlexaFluor 488 azide (Invitrogen, A10266) according to manufacturers’ instructions. Subsequently, Duolink proximity ligation assay (PLA) was performed following the manufacturer’s protocol (MilliporeSigma). The antibodies used in Mira are rabbit anti-biotin (Cell Signaling Technology, 5597S), and mouse anti-biotin (Sigma, B7653). Cytoplasmic MIRA signals were imaged using the Nikon Eclipse Ti inverted microspore and analyzed using Nikon NIS-elements software.

### Determination of mitochondrial superoxide status (MitoSOX staining)

The status of ROS and oxidative stress in MAF cells were examined by determining mitochondrial superoxide using MitoSOX Red (Invitrogen, M36008) staining. MitoSOX Red selectively targets mitochondria and undergoes oxidation by superoxide to form a stable fluorescent compound which was imaged using the red filter of fluorescent microscope. MAFs were grown in 8-well chamber slides for living cells (Invitrogen, 155411). 10 μM Cells were pretreated with Estrogen overnight or for 1 hour before 16μM MTOX incubation as indicated in the Figs., followed by light activation for 5 minutes and recovery for 1 hour. The cells were washed with PBS and HBSS and incubated with 0.5 μM MitoSOX Red reagent for 15 min at 37°C, protected from light. The cells were gently washed three times with warm HBSS buffer and stained with Hoechst for 5 minutes. Live cell imaging was performed using a Nikon Eclipse Ti-U inverted microscope. and analyzed using Nikon NIS-elements software.

### Statistical analysis

Statistical data analysis was performed using GraphPad software. Significant differences between sample groups for mouse experiments were determined using ANOVA and Statistical analysis for DNA fiber assays was determined using the Mann-Whitney test as indicated in the legends. Statistical significance is indicated with asterisks as follows (*p< 0.05, **p < 0.01, ***p < 0.001, ****p < 0.0001, ns p> 0.05).

### Chrystal structure analysis

AlphaFold 3 is used to generate human RAD51C models in the wild-type and L261/L262 deletion variants. The wild-type RAD51C model is consistent with the Alvi-RAD51C (PDB: 8gja) and human-RAD51C (PDB: 8ouz) structures, with RMSD values of 0.64 and 0.57, respectively, particularly the alpha-helix, which contains amino acids 255-276. Both models are superimposed on the human RAD51-DNA model (PDB: 7ejc) to generate the RAD51C-DNA models.

### Data availability

All data is available in the main text or the Supplementary materials.

