## supplemental figures for "Diet induced mitochondrial DNA replication instability in *Rad51c* mutant mice drives sex-bias in anemia of inflammation"

**Supplemental information**

**Figure S1**

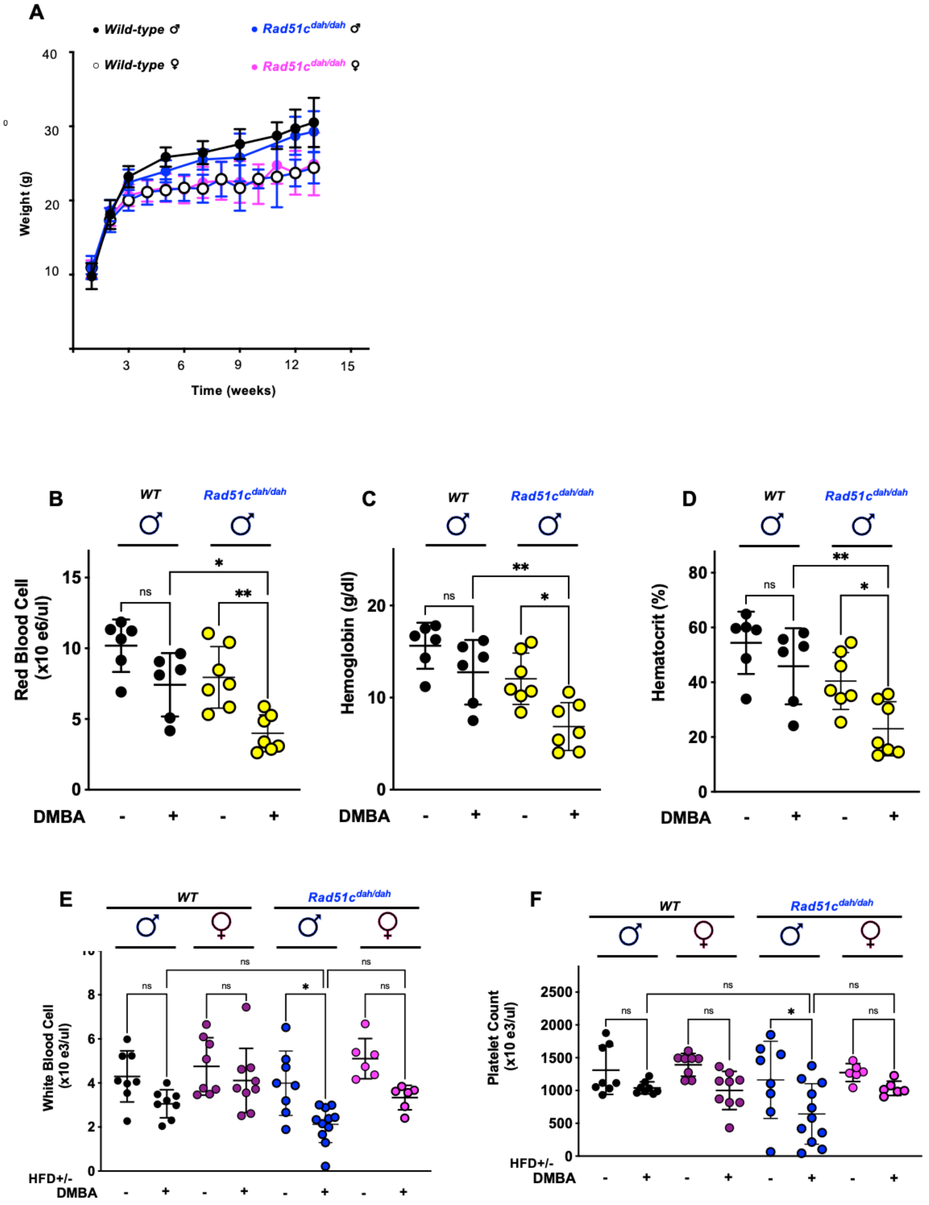


**Fig. S1. Male *Rad51c*$^{dah/dah}$ show pancytopenic features under chronic stress exposure**

**A)** Graph showing weight of Rad51c male and female wild-type (WT) and *Rad51c*$^{dah/dah}$ mice over 14 weeks of age.

**B-D)** Scatter dot plot of red cell blood count (**B**) hemoglobin (**C**) and hematocrit counts (**D**) on normal diet (no HFD) with and without DMBA treatment. Values from Male wild-type (WT) mice are shown in black, and from male *Rad51c*$^{dah/dah}$ mice in yellow.

**E-F)** Scatter dot plot of white blood cell count (**E**) and platelets (**F**) from chronic stress experiments (with HFD) with or without DMBA. Male WT mice (black), female WT mice (purple), male *Rad51c*$^{dah/dah}$ mice (blue) and female *Rad51c*$^{dah/dah}$ (pink).

Horizontal bars represent the mean and error bars represent the SEM. p-values are derived using the one-way Anova test. ns $p>0.05$, * $p<0.05$, **$p < 0.01$, ***$p < 0.001$, ****$p<0.0001$, n=6-11

**Figure S2**

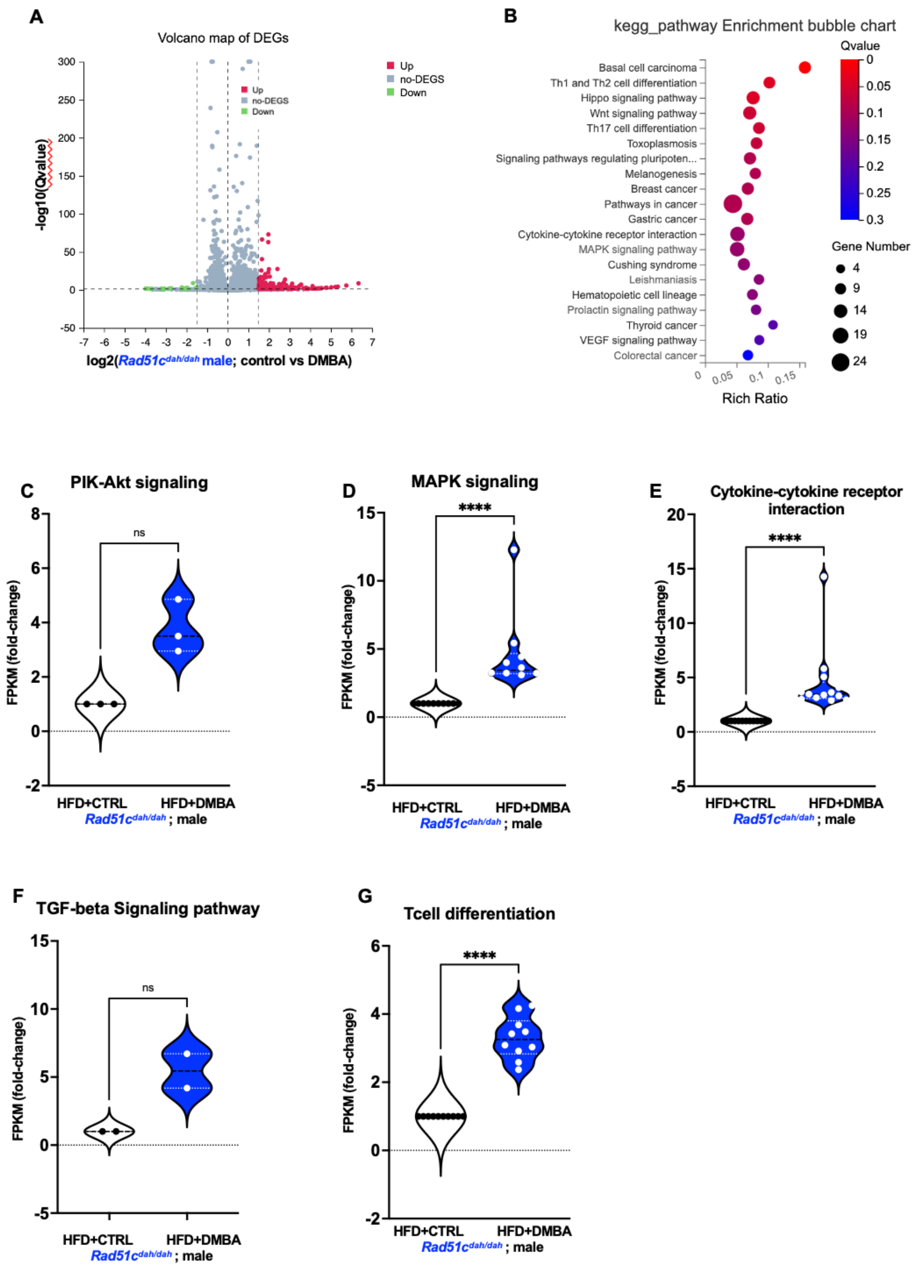


**Fig. S2. Dietary stress induces inflammation in male *Rad51c*$^{dah/dah}$ mice**

**A)** Volcano plot of up- and downregulated genes in bone marrow of male *Rad51c*$^{dah/dah}$ mice with and without DMBA.

**B)** KEGG pathway analysis of upregulated genes in *Rad51c*$^{dah/dah}$ mice with and without DMBA, highlighting upregulation of inflammation pathways.

**C-G)** Box plots showing higher mRNA transcript levels of genes belonging to PI3K-Akt (**C**), MAPK pathway (**D**), cytokine-cytokine receptor interaction pathway (**E**), TGFb-signaling pathway (**F**) and Tcell differentiation pathway (**G**) in male *Rad51c*$^{dah/dah}$ bone marrow with DMBA (right, blue) or without (left, white). Gene set for each pathway (**C-G**) were obtained from gene listed inn KEGG pathway analysis from **B**).

Figure S3

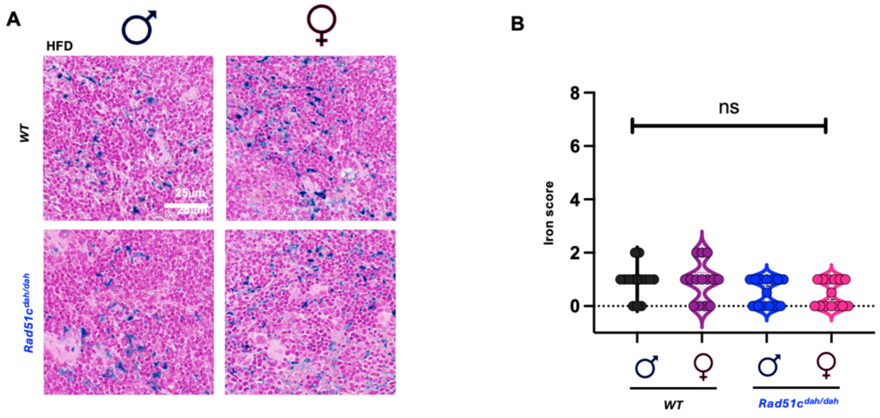


**Fig. S3. Male mice are prone to diet-induced anemia of inflammation**

- **A)** Ferric iron stains of mouse spleen tissue sections of male (left) and female (right) WT (top) and mutant *Rad51c*$^{dah/dah}$ (bottom) mice treated with HFD (no DMBA).
- **B)** Violin plot of (**D**) n=14 image fields for each condition. p-values are derived using the Student T-test. ns p>0.05, * p<0.05, **p < 0.01, ***p < 0.001, ****p<0.0001,

Figure S4

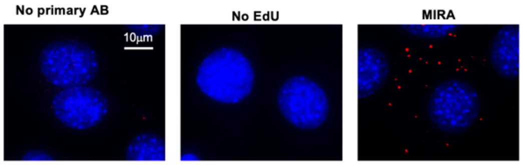


**Fig. S4. MIRA negative controls.**

MIRA (mitochondrial replication assay) signals are unproductive in the absence of a primary antibody (left image), or of EdU (left image), but produce strong MIRA signals in their presence (right image) verifying signal specificity.

Figure S5

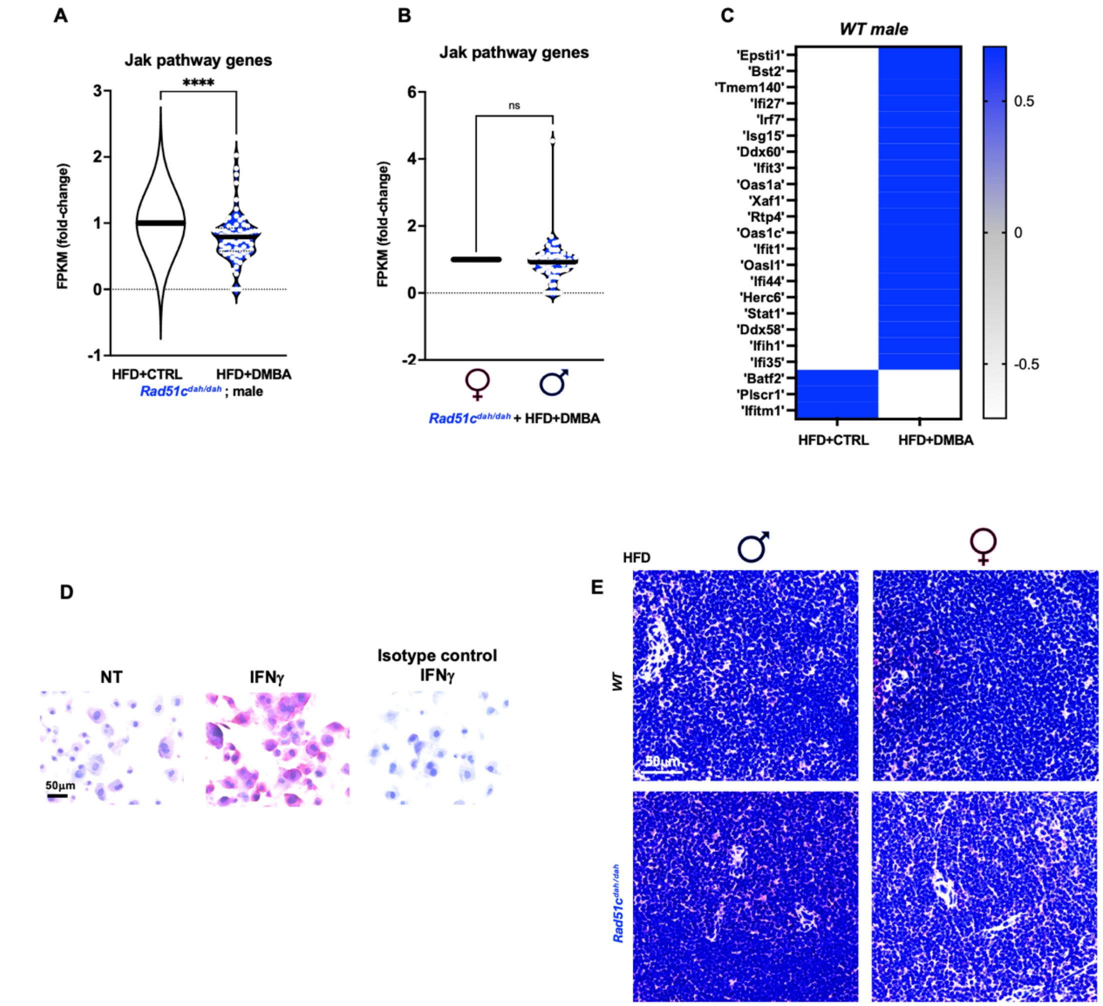


**Fig. S5. Activation of the un-phosphorylated STAT1 response is stronger in male mice**

- **A)** Violin plots of JAK pathway gene expression in male *Rad51c$^{dah/dah}$* mice with and without DMBA
- **B)** Violin plots of JAK pathway gene expression in male and female *Rad51c$^{dah/dah}$* mice with DMBA
- **C)** RNAseq data heatmap of the un-phosphorylated STAT1 response pathway in male high-fat diet-fed wildtype (WT) bone marrow with and without DMBA.
- **D)** Specificity control for STAT1 antibody used for AP red stains. AP red signal is greatly increased in male *Rad51c$^{dah/dah}$* MAFs treated with IFNγ, which induces STAT1, compared to slides that are incubated with an isotype control antibody.
- **E)** Representative images Red Alkaline phosphatase stains against STAT1 in mouse spleen tissue sections (white pulp) of male (left) and female (right) WT (top) and mutant *Rad51c$^{dah/dah}$* (bottom) mice treated with HFD only (without DMBA). Scale bar, 50μm

Figure S6

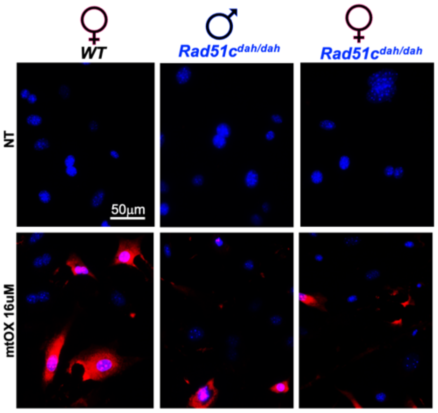


**Fig.. S6 Males are less protected against mitochondrial reactive oxygen species.** Representative images of mitochondrial superoxide staining in male and female wild-type (WT) and female *Rad51c*$^{dah/dah}$ MAF cells using MitoSOX with or without mtOX (16µM) as indicated.

Figure S7

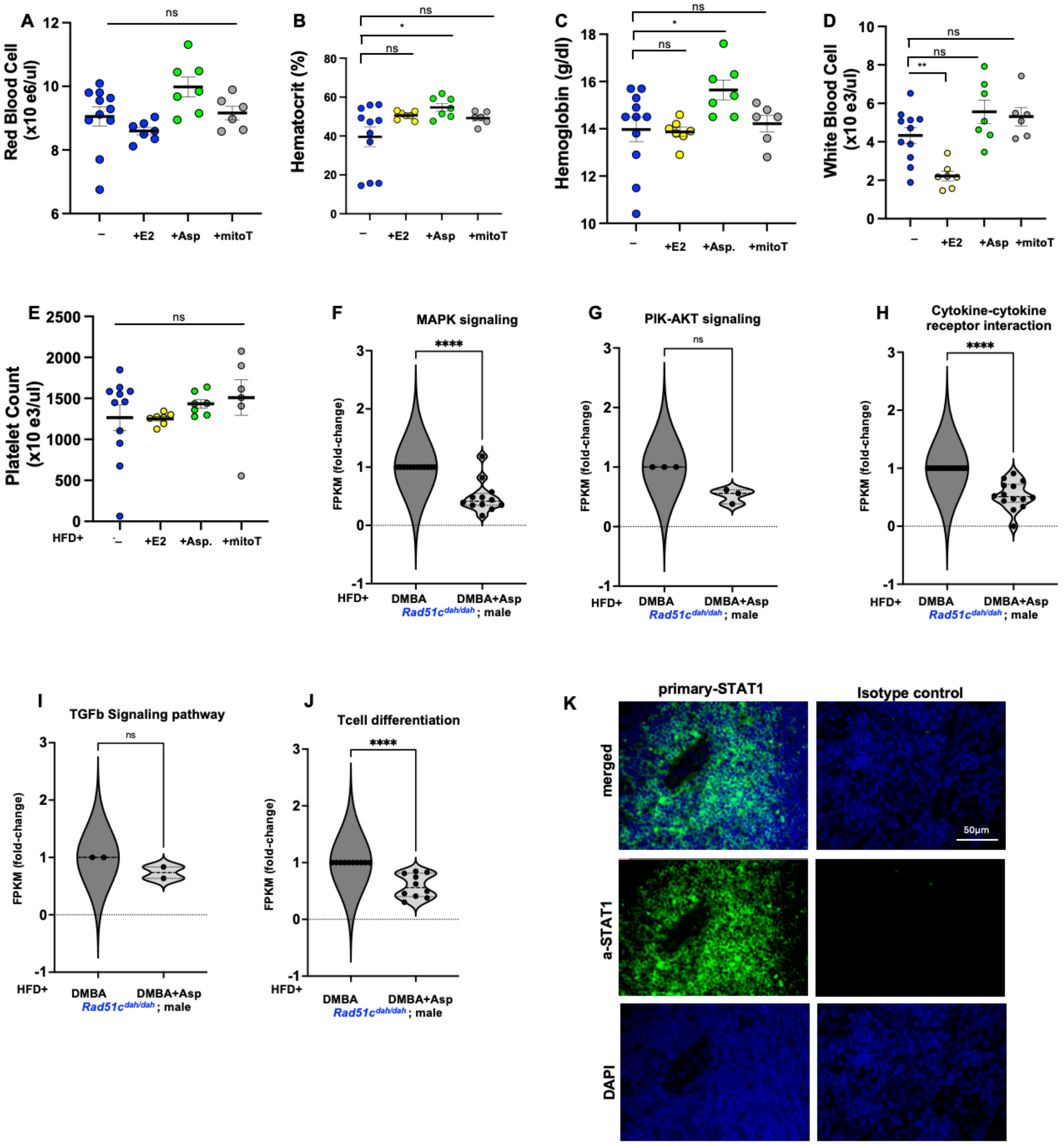


**Fig. S7 Blood counts and macrophage activation pathways by restoring mitochondrial stability under chronic stress exposure**

**A-E)** Red cell blood count (**A**) hematocrit (**B**), hemoglobin (**C**), white blood cell (**D**), and platelet count (B) from male *Rad51c*$^{dah/dah}$ mice without DMBA treatment (blue), treatment with estrogen (yellow), aspirin (green) or mitoTempo (grey).

**F- J)** Violin plots of MAPK signaling pathway gene expression (**F**) PI3K-Akt pathway genes (**G**) cytokine-cytokine receptor interaction pathway genes (**H**) TGFb pathway (**I**) and Tcell differentiation pathway (**J**) for male (blue) *Rad51c*$^{dah/dah}$ mice treated with HFD+DMBA compared to those treated with HFD+DMBA+aspirin (light grey).

**K)** Specificity control for STAT1 antibody used for Immunofluorescence (IF) microscopy. STAT1 IF signal (green) is greatly increased in male *Rad51c*$^{dah/dah}$ HFD+DMBA treated spleen tissue sections(white pulb) compared to slides that are incubated with an isotype control antibody. Blue channel, DAPI (nuclear DNA). Scale bar, 50μm

**Table S1 Composition of high-fat diet and standard diet fed to FVB/NJ mice at 3 weeks**

| Diet | High-fat | Standard |
|---|---|---|
| Protein (% kcal) | 23.1 | 16.9 |
| Carbohydrate (% kcal) | 25.9 | 67.4 |
| Fat (% kcal) | 34.9 | 4.3 |
| Fiber (% kcal) | 6.5 | 4.7 |
| Energy (kcal $g^{-1}$) | 5.1 | 3.76 |
| Ingredients Comosition | % | % |
| Lard | 31.66 | 1.896 |
| Casein | 25.845 | 18,956 |
| Dextrin | --- | 29.856 |
| Maltodextrin | 16.153 | 3.317 |
| Sucrose | 8.847 | 33.129 |
| Cellulose | 6.461 | 4.739 |
| Soybean oil | 3.291 | 2.37 |
| Potassium Citrate | 2.192 | 1.564 |
| Calcium Phosphate | 1.68 | 1.232 |
| Mineral Mix | 1.292 | 0.948 |
| Vitamin Mix | 1.292 | 0.948 |
| Calcium Carbonate | 0.711 | 0.521 |
| L-Cystine | 0.388 | 0.284 |
| Choline Bitartrate | 0.258 | 0.19 |

**Table S2. Birth rates of *Rad51c*$^{WT/dah}$ mating.**

| Parental genotype (♀ & ♂ *Rad51c*$^{WT/dah}$) | | | |
|---|---|---|---|
| Rad51c +/+ | Rad51c +/mt | Rad51c mt/mt | Statistic |
| 26.39%<br>N = 38 | 48.61%<br>N = 70 | 25%<br>N = 36 | $X^2 = 0.167$<br>$p = 0.92$ |

| Table S3 KEGG pathway analysis of 198 upregulated genes in male Rad51c dah/dah DMBA-treated mice compare to females Rad51c dah/dah DMBA-treated. | | | |
|---|---|---|---|
| KEGG pathway Term ID | KEGG Pathway Term | Term Candidate Gene Number | P value |
| 470 | D-Amino acid metabolism | 2 | 0.001031484 |
| 330 | Arginine and proline metabolism | 4 | 0.001075167 |
| 4931 | Insulin resistance | 5 | 0.002299351 |
| 4010 | MAPK signaling pathway | 7 | 0.01172869 |
| 220 | Arginine biosynthesis | 2 | 0.0121046 |
| 5134 | Legionellosis | 3 | 0.01400526 |
| 4066 | HIF-1 signaling pathway | 4 | 0.0155339 |
| 4936 | Alcoholic liver disease | 4 | 0.03033348 |
| 4060 | Cytokine-cytokine receptor interaction | 6 | 0.03577368 |
| 5226 | Gastric cancer | 4 | 0.03689551 |
| 4657 | IL-17 signaling pathway | 3 | 0.04135366 |
| 250 | Alanine, aspartate and glutamate metabolism | 2 | 0.04260485 |
| 620 | Pyruvate metabolism | 2 | 0.05295387 |
| 4933 | AGE-RAGE signaling pathway in diabetic complications | 3 | 0.05347959 |
| 4144 | Endocytosis | 5 | 0.0773618 |
| 4151 | PI3K-Akt signaling pathway | 6 | 0.08188126 |
| 4611 | Platelet activation | 3 | 0.08697867 |
| 4213 | Longevity regulating pathway - multiple species | 2 | 0.09619119 |
| 5217 | Basal cell carcinoma | 2 | 0.0988216 |
| 4915 | Estrogen signaling pathway | 3 | 0.1037518 |

**Table S3 KEGG pathway analysis of 198 upregulated genes in male Rad51c dah/dah DMBA-treated mice compare to females Rad51c dah/dah DMBA-treated.**

| KEGG pathway Term ID | KEGG Pathway Term | Term Candidate Gene Number | P value |
|---|---|---|---|
| 470 | D-Amino acid metabolism | 2 | 0.001031484 |
| 330 | Arginine and proline metabolism | 4 | 0.001075167 |
| 4931 | Insulin resistance | 5 | 0.002299351 |
| 4010 | MAPK signaling pathway | 7 | 0.01172869 |
| 220 | Arginine biosynthesis | 2 | 0.0121046 |
| 5134 | Legionellosis | 3 | 0.01400526 |
| 4066 | HIF-1 signaling pathway | 4 | 0.0155339 |
| 4936 | Alcoholic liver disease | 4 | 0.03033348 |
| 4060 | Cytokine-cytokine receptor interaction | 6 | 0.03577368 |
| 5226 | Gastric cancer | 4 | 0.03689551 |
| 4657 | IL-17 signaling pathway | 3 | 0.04135366 |
| 250 | Alanine, aspartate and glutamate metabolism | 2 | 0.04260485 |
| 620 | Pyruvate metabolism | 2 | 0.05295387 |
| 4933 | AGE-RAGE signaling pathway in diabetic complications | 3 | 0.05347959 |
| 4144 | Endocytosis | 5 | 0.0773618 |
| 4151 | PI3K-Akt signaling pathway | 6 | 0.08188126 |
| 4611 | Platelet activation | 3 | 0.08697867 |
| 4213 | Longevity regulating pathway - multiple species | 2 | 0.09619119 |
| 5217 | Basal cell carcinoma | 2 | 0.0988216 |
| 4915 | Estrogen signaling pathway | 3 | 0.1037518 |

| Table S4 KEGG pathway analysis of 572 upregulated genes in male Rad51c dah/dah DMBA-treated mice compare to male Rad51c dah/dah control | | | |
|---|---|---|---|
| KEGG pathway Term ID | KEGG Pathway Term | Term Candidate Gene Number | P value |
| 5217 | Basal cell carcinoma | 10 | 2.41E-06 |
| 4658 | Th1 and Th2 cell differentiation | 9 | 2.70E-04 |
| 4390 | Hippo signaling pathway | 12 | 4.28E-04 |
| 4310 | Wnt signaling pathway | 12 | 8.28E-04 |
| 4659 | Th17 cell differentiation | 9 | 9.95E-04 |
| 5145 | Toxoplasmosis | 9 | 0.00138452 |
| 4550 | Signaling pathways regulating pluripotency of stem cells | 10 | 0.00213175 |
| 4916 | Melanogenesis | 8 | 0.00291859 |
| 5224 | Breast cancer | 10 | 0.00304728 |
| 5200 | Pathways in cancer | 24 | 0.00306542 |
| 5226 | Gastric cancer | 10 | 0.00335962 |
| 4060 | Cytokine-cytokine receptor interaction | 15 | 0.00482299 |
| 4010 | MAPK signaling pathway | 15 | 0.00529691 |
| 4934 | Cushing syndrome | 10 | 0.00606151 |
| 5140 | Leishmaniasis | 6 | 0.00689869 |
| 4640 | Hematopoietic cell lineage | 7 | 0.0073383 |
| 4917 | Prolactin signaling pathway | 6 | 0.00900037 |
| 5216 | Thyroid cancer | 4 | 0.01194603 |
| 4370 | VEGF signaling pathway | 5 | 0.01303487 |
| 5210 | Colorectal cancer | 6 | 0.01994574 |
